# Endophytic bacterial communities of alpine Rosaceae plants are affected by the plant tissue, collection site and host plant and culturable psychrotolerant isolates contribute to plant freezing stress tolerance

**DOI:** 10.1101/2023.11.18.567389

**Authors:** Malek Marian, Livio Antonielli, Ilaria Pertot, Michele Perazzolli

## Abstract

- Wild plants growing in alpine regions are associated with endophytic microbial communities that may support plant growth and survival under cold conditions.
- The structure and function of endophytic bacterial communities were characterised in flowers, leaves and roots of three alpine Rosaceous plants in Alpine areas using a combined amplicon sequencing and culture-dependent approach to identify factors shaping these communities.
- Amplicon-sequencing analysis revealed that plant tissue, collection site and host plant are the main factors affecting the richness, diversity and taxonomic structure of endophytic bacterial communities in alpine Rosaceae plants. Core endophytic bacterial taxa were identified as 31 amplicon sequence variants highly prevalent across all plant tissues.
- Psychrotolerant bacterial endophytes belonging to the core taxa of *Duganella, Erwinia, Pseudomonas* and *Rhizobium* genera mitigated freezing stress in strawberry plants, demonstrating the beneficial role of endophytic bacterial communities and their potential use for cold stress mitigation in agriculture.

## Introduction

Plants are associated with highly diverse, complex and dynamic microbial communities, which are hosted on (i.e. epiphytic) and inside (i.e. endophytic) plant tissues and can support plant growth under stress conditions (Compant *et al*., 2019). In particular, endophytic bacterial communities have intimate interactions and extensive exchange of metabolites with their plant host (Compant *et al*., 2021). Plant-associated microbial communities have complex and diverse structures, and identifying which members are functionally important for the host plant is a still a key challenge in plant microbiome research. A small subset of plant-associated microbial communities constitutes the core microbial taxa and consists of a group of members present in almost all the communities associated with a given host plant across a wide range of environments with potential beneficial traits for the plant host (Trivedi *et al*., 2020). For example, core microbial taxa play important roles in promoting plant growth and health by fixing nitrogen (Zhang *et al*., 2022a) and/or suppressing plant diseases (Hong *et al*., 2023). The core microbial taxa may also include the so-called ‘hub taxa’ (or keystone taxa), which can be defined as highly connected nodes in co-occurrence network analysis (Agler *et al*., 2016; Banerjee *et al*., 2018). Hub taxa can drive microbial community structure and support functions, for example they can support soil microbial-mediated functions (e.g. nutrient cycling, organic matter decomposition and gas emissions) in the presence of a loss of microbial biodiversity (Luo *et al*., 2023) or mediate microbial extracellular enzyme degradation and assimilation (Wang *et al*., 2023). Therefore, the identification of functional hub taxa in the core microbiota of plants could be the first step to identify the key microbial community members to be further manipulated to improve plant stress tolerance.

Wild plants growing in alpine regions are associated with cold-adapted, highly complex and host-specific microbial communities, which may promote plant growth and survival under cold conditions (Marian *et al*., 2022). For example, psychrotolerant (i.e. cold adapted) *Pseudomonas* spp. isolated from the leaf apoplast of five mountain plant species (*Colchicum speciousum*, *Draba nemorosa*, *Erodium cicutarium*, *Galanthus gracilis* and *Scilla siberica*) improved bean (*Phaseolus vulgaris* L.) tolerance to freezing stress (Tiryaki *et al*., 2019). Likewise, *Paraburkholderia phytofirmans* PsJN can mitigate freezing damage in *Arabidopsis thaliana* possibly by inducing cell wall strengthening (Su *et al*., 2015).

Fruit crops belonging to the Rosaceae family, such as apple, cherry, pear and strawberry, are of relevant economic and agricultural value in temperate regions (Shulaev *et al*., 2008). Unfortunately, freezing stress is one of the major abiotic stresses affecting fruit production in Rosaceae crops, causing damage to flower and leaf tissues in the spring period (Rodrigo, 2000; Salazar-Gutiérrez *et al*., 2016). On average, spring frost occurs at least once every three to five years and causes severe losses in many temperate regions in the European Union, United States and China (Wolfe *et al*., 2018; Vitasse & Rebetez 2018; Ru *et al*., 2023). Furthermore, global warming due to climate change will likely promote early spring-related phenological events in plants with a consequent increased probability of incurring severe spring frosts (Gu *et al*., 2008; Zohner *et al*., 2020; Ru *et al*., 2023). No information is currently available on the structure and function of endophytic bacterial communities associated with alpine Rosaceous plants (particularly reproductive organs such as flowers) and factors that shape the community structure.

Current strategies implemented by growers to reduce freezing damage consist of physical methods (e.g. overhead irrigation, heaters and wind machines) and chemical treatments (e.g. growth regulators, vitamins, and bactericides) (Román-Figueroa *et al*., 2021; Drepper *et al*., 2022). However, these approaches have several limitations in terms of costs, efficacy, feasibility and environmental impacts (Albrecht *et al*., 2004). The use or manipulation of plant-associated microbial communities was proposed as a promising sustainable approach to alleviate cold stress in crops (Beirinckx *et al*., 2020; Erlandson *et al*., 2021; Persyn *et al*., 2022), but no information is available on the possible mitigation of freezing stress in Rosaceae plants. The aim of this study was to assess the relationship between plant tissue, plant host and collection site and the taxonomic structure of endophytic bacterial communities associated with alpine Rosaceae plants and to identify core taxa and hub taxa using a culture-independent approach. Moreover, culturable psychrotolerant bacterial endophytes were isolated from the same alpine Rosaceae plants to clarify their possible contribution to the mitigation of freezing stress in a model plant (strawberry).

## Materials and Methods

### Plant material and experimental design

Samples of three alpine Rosaceae plants (*Alchemilla* sp., *Dryas octopetala* and *Geum montanum*) were collected in Alpine areas of the Trentino-alto Adige Region, Italy, specifically in seven sites (Val di Non, Val di Sole, Val di Pejo, Val Rendena, South Tyrol, Stelvio Park and Val di Fassa, hereafter named Site A, B, C, D, E, F and G, respectively) and two expositions (North and South) (Fig. S1), to obtain alpine Rosaceae plants from different altitudes and climatic conditions (Table S1). The sampling was carried out from the middle of June to the middle of August 2021 to match the flowering period of each species at each site. For each alpine Rosaceae plant/tissue/site/exposition, three replicates were collected, each of which was collected from 15–20 randomly chosen plants with no visible signs of damage or disease, transported to the laboratory in a cool box, stored in a refrigerator at 4°C and processed within 24 hours after collection. A total of 270 samples (Table S1) were analysed from three alpine Rosaceae plants: 108 samples from *Alchemilla* sp., 54 samples from *D. octopetala* and 108 samples from *G. montanum*.

### Sample processing

Flowers, leaves and roots were cut from each sampled plant and surface disinfected with 70% ethanol for 1 min, 1% sodium hypochlorite for 1 min (flowers), 2.5% sodium hypochlorite for 5 min (leaves), or 9% sodium hypochlorite for 10 min (roots), followed by 70% ethanol for 1 min and five washes with sterile distilled water (SDW) of 2 min each. Samples were air-dried under laminar flow for 30 min. Half of each sample was frozen with liquid nitrogen and stored at −80°C for DNA extraction and amplicon sequencing analysis, whereas the other half was used for the isolation of culturable bacteria. As a control for plant surface disinfection, the last washing solution (20 mL) was centrifuged (3,500 g for 10 min), the supernatant was discarded and aliquots (20 μL) of the remaining solution (500 μL) were plated on solid Reasoner’s 2A (R2A; Sigma-Aldrich, Merck, Rahway, NJ, USA) to confirm the absence of bacterial growth seven days after incubation at 25°C.

### DNA extraction, amplification, library preparation and amplicon sequencing

Culture-independent analysis of DNA extraction, amplification and sequencing was carried out as previously described with some modifications (Perazzolli *et al*., 2022). Briefly, genomic DNA was extracted from surface-disinfected tissues using the FastDNA Spin Kit for Soil (MP Biomedical, Santa Ana, CA, USA) with a slight modification. In particular, surface-disinfected tissues (1 g) were homogenized in sterile stainless jars on liquid nitrogen using a mixer-mill disruptor (MM 400, Retsch, Haan, Germany) at 25 Hz for 10 sec. Powdered frozen tissues were added to a lysing matric E tube (MP Biomedical) and homogenized with the FastPrep-24 Classic instrument (MP Biomedical) at a speed of 4.0 for 5 sec (flowers) or 30 sec (leaves and roots). The bacterial V5–V7 region of 16S ribosomal DNA (rDNA) was amplified with a nested PCR approach (Mitter *et al*., 2017). The first bacterial 16S amplification was carried out with the primers 799 forward (5’-AACMGGATTAGATACCCKG-3’) and 1392 reverse (5’-ACGGGCGGTGTGTRC-3’), to exclude chloroplast 16S rDNA and to amplify bacterial and mitochondrial rRNA of 600 bp and 1,000 bp amplicon size, respectively (Mitter *et al*., 2017). Bacterial 16S amplicons were purified by agarose gel separation, followed by the NucleoSpin Gel and PCR Clean-up purification kit (Macherey-Nagel, Düren, Germany). The second 16S amplification was performed with the primers 799 forward (5’-AACMGGA TTA GAT ACC CKG-3’) and 1175 reverse (5’-ACGTCRTCCCCDCCTTCCT-3’) including the specific overhang Illumina adapters (5’-ACACTCTTTCCCTACACGACGCTCTTCCGATCT-3’ and 5’-GACTGGAGTTCAGACGTGTGCTCTTCCGATCT-3’, respectively) for amplicon library construction, and 16S amplicons (500 bp amplicon size) were purified by agarose gel separation using the NucleoSpin Gel and PCR Clean-up purification kit (Macherey-Nagel). Bacterial 16S amplifications were obtained using the FastStart High-Fidelity PCR system (Roche, Branford, CT, USA) with PCR amplification conditions optimised for each plant tissue (Table S2). A quality check was performed by gel electrophoresis to confirm the presence of the expected amplicon using 5 µL of PCR product. PCR product purification, quantification, library construction and sequencing were conducted by Eurofins Genomics GmbH (Ebersberg, Germany) following their in-house NGSelect Amplicons approach with an index PCR to introduce indexed sequencing adaptors for sample discrimination after Illumina MiSeq (PE300 mode) sequencing.

### 16S rRNA gene amplicon sequence processing

16S rRNA gene amplicon sequence processing was carried out as previously described (Perazzolli *et al*., 2022). Briefly, Illumina reads were filtered with Bowtie2 v2.4.2 (Langmead & Salzberg, 2012), sequence quality was checked with FastQC v0.11.9 and primers were cut using Cutadapt v3.4 (Martin, 2011). Sequences were quality filtered, trimmed, denoised (filtered read counts) and ASVs were generated with DADA2 v1.18.0 (Callahan *et al*., 2016). Denoised forward and reverse ASV sequences were merged and chimera filtered. Bacterial ASVs were checked using Metaxa2 v2.2.3 (Bengtsson-Palme *et al*., 2016) for targeting the presence of the V5 to V7 hypervariable regions of the 16S rRNA gene. Taxonomy was assigned using the RDP classifier implemented in DADA2 against the SILVA v138.1 (Quast *et al*., 2013). A bacterial table of read counts was built and imported into the R-4.3.0 statistical environment for further analyses (R Core Team, 2023). After taxonomic classification, ASVs classified as plastid rRNA and other than archaea or bacteria were removed. ASVs with no reads or singletons, as well as very low-abundance ASVs with a maximum relative abundance below 0.1% per sample, were discarded before any further analysis.

### Analyses of 16S rRNA gene amplicon sequence data

Bacterial ASV tables were split into two datasets: (i) dataset for *Alchemilla* sp. and *G. montanum* from six collection sites (hereafter termed dataset 1; 216 samples) and (ii) dataset for *Alchemilla* sp., *D. octopetala* and *G. montanum* from two collection sites (hereafter termed dataset 2; 108 samples). This approach was performed to obtain consistent numbers of alpine Rosaceae plants from the different collection sites since *D. octopetala* was not present in some collection sites (Table S1). In particular, the design incorporated four categorical orthogonal factors in dataset 1, such as Tissue (three levels: flowers, leaves and roots), alpine Rosaceae plants (two levels: *Alchemilla* and *Geum*), Collection site (six levels: Site A, B, C, D, F and G) and Exposition (two levels: North and South), and four categorical orthogonal factors in dataset 2, such as Tissue (three levels: flowers, leaves and roots), alpine Rosaceae plants (three levels: *Alchemilla*, *Dryas* and *Geum*), Collection sites (two levels: Site D and G) and Exposition (two levels: North and South).

Alpha-diversity values were calculated by the multiple rarefaction method for both richness (observed ASVs) and diversity (estimated with Simpson’s index) values by averaging the results inferred after 999 rarefactions, starting with the lowest read counts in a sample (*n* = 13,869) for the complete dataset, using the rtk R package (Saary *et al*., 2017). The rarefaction was repeated for each dataset and then values were fitted to the linear models (LMs) together with the four factors (tissue, alpine Rosaceae plant, collection site, and exposition). LMs with the lowest root mean squared error (RMSE) identified using the caret R package (Kuhn, 2008) were chosen for the analysis of variance (ANOVA) followed by post-hoc analysis with estimated marginal mean comparisons using the emmeans R package (Lenth, 2023). Conditional inference regression tree analysis was further applied to visualize the hierarchy among different factors on the alpha-diversity metrics (Hothorn & Zeileis, 2021).

Beta-diversity values were normalised using the multiple rarefaction method to account for differences in sequencing depth as described above for the alpha-diversity. Permutational multivariate analysis of variances (PERMANOVA) global test and PERMANOVA partitioning test by tissues were conducted on Bray–Curtis dissimilarity matrices using the adonis2 function from the vegan R package (Oksanen *et al*., 2023) to determine the differences in centroids of the bacterial communities across samples in dataset 1 and dataset 2. In addition, intra-factor pairwise comparisons between levels were carried out using pairwise.perm.manova function from the RVAideMemoire R package with *P*-value adjustment using Benjamini-Hochberg method (Herve, 2023). To identify the drivers of bacterial community structure in each tissue, PERMANOVA tests were performed separately for the different tissues. In addition, the host-environment effects index (HEEI = relative contribution of alpine Rosaceae plants/relative contribution of the collection site) was calculated based on PERMANOVA (Xiong *et al*., 2021). A constrained analysis of principal coordinates (CAP) based on Bray–Curtis dissimilarity matrices was carried out using the capscale function from the vegan package, and the significance of constraints was confirmed by a permutational test (999 iterations). Differences in multivariate homogeneity of group dispersions (PERMIDISP2) were also evaluated using the betadisper function from the vegan package, followed by ANOVA and permutational test (999 iterations). The distance-based PERMANOVA was complemented with multivariate generalized linear models (mGLMs), which account for the mean-variance relationship of the data (Wang *et al*., 2012; Warton *et al*., 2012). mGLMs were fitted to the rarefied counts of bacterial ASVs with an occupancy (frequency of detection) of 0.25 to reduce complexity and computation time. A negative binomial distribution of the data was graphically verified by plotting fitted *vs* residual values, as previously described (Bálint *et al*., 2015). The manyglm function from the mvabund R package (Wang *et al*., 2012) was used with the full model (Tissue*alpine_Rosaceae_plant*Collection_site*Exposition), as it was identified to be the best-performing model based on the obtained Akaike information criterion (AIC) values. An analysis of deviance (Dev) was calculated with a likelihood-ratio test using a permutational test (1000 iterations, Monte Carlo resampling). A model-based ordination diagram was generated using the ordiplot function after performing generalised linear latent variable models with the variables using the gllvm R package (Niku *et al*., 2019). The model fit was restricted to five latent variables as identified by the lowest AIC values.

Prior to hierarchical clustering, rarefied count data were standardised with the Wisconsin double standardization method, and Bray–Curtis dissimilarity matrices were calculated using the vegdist function from the vegan package. Hierarchical clustering analysis was performed using the agnes function with the Ward. D method (identified as the strongest clustering structure by agglomerative coefficient test) from the cluster R package (Maechler *et al*., 2022).

Core endophytic bacterial taxa of Rosaceae plants (set of bacterial taxa that are characteristic of a host plant; Trivedi *et al*., 2020) were identified based on the abundance and occupancy distribution as previously described (Shade & Stopnisek, 2019, Stopnisek & Shade, 2021). Briefly, the mean relative abundance and occupancy of each taxon were calculated across the rarefied count data of the complete dataset. Taxa were then ranked by occupancy with an additional weight for taxa with an occupancy of 1 in a particular tissue. Bray–Curtis dissimilarities were calculated for the complete dataset starting from the first ranked taxa. The final 2% increase in the collective contribution of ranked taxa to the Bray–Curtis dissimilarities were used to infer core taxa members. Taxa abundance/occupancy data were further fitted to the Sloan neutral model [95% confidence intervals, goodness of fit (R2) and estimated migration rate (*m*; a measure of dispersal limitation)] to evaluate the importance of stochastic and deterministic processes in the overall bacterial community assembly. In addition, the relative importance of each assembly process of homogeneous and heterogeneous selection (i.e. deterministic), as well as dispersal limitation, homogenizing dispersal, drift and others (i.e. stochastic) was investigated using the NST and iCAMP (Infer Community Assembly Mechanisms by Phylogenetic-bin-based null model) R packages (Ning *et al*., 2019; 2020). Briefly, the dominance of deterministic and stochastic assemblies was determined with NST when the phylogenetic normalized stochasticity ratio (pNST) was lower and higher than 50%, respectively. The parameter settings for iCAMP were 0.2 phylogenetic distance, 24 minimal bin size requirement threshold and SES.RC as the null model significance test. The latter setting uses the beta net relatedness index (βNRI) for phylogenetic beta diversity and the modified RaupCrick for taxonomic beta-diversity. For the community assembly analysis, a phylogenetic tree was constructed by aligning all ASVs and calculating pairwise distance between them before tree inference using the msa, seqinr and ape R packages (Bodenhofer *et al*., 2015; Charif & Lobry 2007).

Random forest models were performed to identify taxa with strong associations with the four factors (i.e. tissue, alpine Rosaceae plant, collection site and exposition) using a machine learning algorithm, as previously described (Perazzolli *et al*., 2022). Rarefied count data were used as predictors, and factors were used as response data. Model performances were evaluated with repeated k-fold cross-validation (tenfold, 10 repetitions) and mtry values that determined the highest model accuracy were chosen and input to Random Forest analysis, as implemented in the caret R package (Kuhn, 2008). Variable importance was assessed with permutations (999 iterations) using the rfPermute R package. ASVs with significant mean decrease accuracy (*P* ≤ 0.05) in all four factors were extracted (selected ASVs) and used for the differential abundance analysis. To this end, the analysis of compositions of microbiomes with bias correction 2 (ANCOM-BC2) was used on nonrarefied count data (Lin & Peddada, 2020). ASVs with structural zeros (i.e. ASVs present in at least one group of flowers, leaves and roots but absent in at least one group of other tissues) were not considered in further data analysis. Plant tissue, alpine Rosaceae plant, collection site and exposition were implemented as four covariates in the model formula (Tissue+alpine_Rosaceae_plant+Collection_site+Exposition), and ANCOM-BC2 primary analysis was used to determine ASVs that were differentially abundant according to covariates. All log-ratios for each taxon were then tested for significance using a two-sided Z-test with W, and *P*-values were adjusted using the Benjamini‒Hochberg method (Lin & Peddada, 2020).

Co-occurrence networks were constructed to characterize the relationship between core taxa and the whole community using the NetCoMi v1.1.0 package, as previously described (Olofintila & Noel, 2023; Peschel *et al*., 2021), and hub ASVs were identified as those that are significantly more connected within the network than other ASVs on the basis of centrality measures (Banerjee *et al*., 2018). To construct the correlation matrix, the SpiecEasi v1.1.2 tool (Kurtz *et al*., 2015) was used on nonrarefied count data as SpiecEasi performs centered log-ratio normalisation internally. The data consisted of ASVs with 0.001% relative abundance and occupancy of 0.25 for each group of tissue (i.e. flowers, leaves and roots). Association matrices were estimated using the Meinshausen and Bühlmann algorithm with the nlambda set to 100, sampled 100 times, lambda.min.ratio set to 10^−1^, and with the stars model selection. Matrices were then used to construct the network using the netConstruct function and network properties were determined using the netAnalyze function. Hub nodes were identified based on eigenvector centrality values above 95% of the empirical distribution of all eigenvector centralities in the network (Peschel *et al*., 2021). The network was visualized using the plot.microNetProps function. Quantitative network comparisons were performed with the netCompare function using 1000 permutations, and the Jaccard index was used to assess how different the sets of most central nodes were between the two groups (0 if the sets were completely different and 1 for exactly equal sets).

Multiple sequence alignment for both Sanger (psychrotolerant bacterial endophytes) and ASVs (most abundant bacterial ASV, mean relative abundance > 0.1%) sequences was performed using MAFFT v7 (Katoh *et al*., 2019). A phylogenetic tree was inferred from partial 16S rRNA gene sequences based on the neighbor-joining algorithm with the Jukes-Cantor correction and 1000 resampling bootstraps (Katoh *et al*., 2019). The tree was visualized and annotated using Archaeopteryx and iTOL v5, respectively (Han & Zmasek, 2009; Letunic & Bork, 2021).

All plots were generated using the data visualization R packages ggplot2 v3.4.2 (Wickham, 2016) and patchwork v1.1.2 (Pedersen, 2023).

### Isolation and taxonomic annotation of culturable psychrotolerant bacterial endophytes

Isolation and identification of the bacterial endophytes of alpine Rosaceae plants were performed according to previously described protocols (Arrigoni *et al*., 2018; Nissinen *et al*., 2012). In brief, surface-disinfected flower, leaf or root tissues (1 g) originating from the same plant samples used for amplicon sequencing were homogenised in SDW using refrigerated (ice) sterile stainless steel with a mixer-mill disruptor (MM 400, Retsch) at 25 Hz for 30 sec, 2 min, or 10 min for flowers, leaves and roots, respectively. Culturable bacteria were isolated by plating serial dilutions of each suspension (100 µL aliquots) on solid R2A medium supplemented with 50 mg L^−1^ cycloheximide to prevent fungal contamination. Plates were incubated at 4°C to allow the growth of psychrotolerant isolates, and bacterial colony forming units (CFU) per unit of plant fresh weight (CFU g^−1^) were assessed after 30 days. Three technical replicate plates were used for each dilution and each sample.

Representative bacterial isolates were selected for each sample based on morphological visual observation of bacterial colonies (e.g. size, colour, opacity, texture, form, elevation and margin), as previously described (Perazzolli *et al*., 2022). The purified colonies were suspended in 40% (v/v) glycerol and kept at −80°C until use. To avoid redundancy, bacterial isolates were taxonomically annotated based on sequencing of the V6–V8 region of the 16S rRNA gene. PCR amplification and DNA sequencing were performed with the universal primers 27 forward (5’-AGAGTTTGATCCTGGCTCAG-3’ and 1492 reverse (5’-GGTTACCTTGTTACGACTT-3’) using the conditions previously described (Perazzolli *et al*., 2022). Briefly, the PCR product was purified using a NucleoSpin Gel and PCR Clean-up purification kit (Macherey-Nagel) and subjected to Sanger sequencing at Eurofins Genomics GmbH (Ebersberg, Germany). Forward and reverse sequences were assembled by using Chromas Pro software version 1.7 (Technelysium Pty Ltd., Tewantin, Australia), and taxonomy was assigned to the genus level by comparing sequences with those of the 16S ribosomal RNA database in the National Center for Biotechnology Information (NCBI) using the rBLAST and taxonomizr R packages (Hahsler & Nagar, 2019; Sherrill-Mix, 2023). To obtain a list of representative psychrotolerant bacterial endophytes, a custom-built database containing all the recovered ASV sequences (*n* = 35,300 ASVs before any filtering) was constructed using the makeblastdb function from the rBLAST package. Sanger sequences were then queried against the customized ASV database using BLAST, parameters of identity 95%, E-value 1 × 10^-5^, max target 10 and max HSPS 10 were used with the predict function of the rBLAST package and taxa of culturable psychrotolerant bacterial endophytes corresponding to the most abundant ASVs (mean relative abundance > 0.1%) were considered as representative psychrotolerant bacterial endophytes.

### Screening of psychrotolerant bacterial endophytes for their freezing protection ability

Representative psychrotolerant bacterial endophytes were tested for their freezing protection ability. Each isolate was grown overnight (18 h) in liquid R2A medium at 25°C under orbital shaking at 200 rpm and bacterial cells were collected by centrifugation (3,500 × g for 10 min) and washing (three times) with sterile 10 mM MgSO_4_. The bacterial suspension was then adjusted to an OD_600_ = 0.1 corresponding to approximately1.0 × 10^8^ CFU mL^−1^.

Seeds of strawberry (*Fragaria* × *ananassa*) cultivar Fresca (Moles Seeds Ltd, Essex, United Kingdom) were germinated as previously described (Ito *et al*., 2011). Briefly, seeds were stratified at 4°C for 3 weeks and scarified with cold 96% sulfuric acid (H_2_SO_4_) for 10 min, followed by five washes with SDW (2 min each). Seeds were surface-disinfected with 70% ethanol for 1 min, 1% sodium hypochlorite for 1.5 min, and 70% ethanol for 1 min, followed by five washes with SDW. Surface-disinfected seeds were sown on two layers of filter paper (Whatman, GE Healthcare Life Sciences) in Petri dishes (90 mm in diameter) moistened with 5 mL of SDW. The plates were incubated in a growth chamber at 23°C ± 1°C for four days with a photoperiod of 14 h/10 h light/dark.

Germinated seeds were transferred to 12-well plates (Greiner-Bio one, Merck) containing solid (7 g L^−1^ agar) full-strength Hoagland (2.75 mL well^−1^; pH adjusted to 6.5) (Sigma-Aldrich, Merck), sealed with parafilm and placed under the growth chamber. After 1 week, seedlings were treated with 10 µL of sterile 10 mM MgSO_4_ (mock-inoculated) or inoculated with a bacterial suspension (bacterium-inoculated) by distributing small droplets on the first two leaves (Vogel *et al*., 2021). As an additional control, seedlings were inoculated with *Pseudomonas syringae* B301D (obtained from The Leibniz Institute DSMZ, Germany), as this strain possesses an *ina* gene and is thus capable of accelerating freezing stress damage to plants (Proebsting & Gross 1988). Seedlings were incubated in the growth chamber for 25 days and exposed to freezing stress as previously described (Jiang *et al*., 2020). Briefly, plates were placed in a freezing chamber (MIR-154, Sanyo, Japan) at 4°C for 30 min, and the temperature was adjusted by a cooling rate of 2°C h^−1^ and then held at −6°C for 6 h in darkness. After the freezing treatment, seedlings were allowed to thaw in darkness for 4 h by increasing the temperature up to 4°C at a rate of 2°C h^−1^ before being transferred back to the growth chamber for 24 h.

Electrolyte leakage was measured by submerging the seedlings in 15-mL tubes containing 10 mL of SDW. Tubes were shaken overnight (18 h) at 150 rpm at 25°C, and the electrical conductivity (EC1) was measured using a hand-held conductivity meter (pH CO 1030, pH/Conductivity Tester VWR, Milan, Italy). Samples were autoclaved and the total electrical conductivity (EC2) was measured. Electrolyte leakage was calculated as follows: (EC1)/(EC2) × 100. The reduction in electrolyte leakage (%) was calculated as follows: [1 − (mean percent electrolyte leakage in the bacterial treatment)]/ (mean percent electrolyte leakage in the control treatment) × 100. In the first trial, six replicates (seedlings) were analysed for each representative psychrotolerant bacterial endophyte, and the best performing isolates were selected. To validate the results, six replicates (seedlings) were analysed for each best performing isolate of representative psychrotolerant bacterial endophytes, and the experiment was carried out three times.

At the end of the freezing bioassay, the colonization of strawberry seedlings by psychrotolerant bacteria was evaluated according to Galambos *et al*. (2020). Briefly, leaf tissues from three strawberry seedlings (0.5 g) were homogenized in sterile stainless jars using a mixer-mill disruptor at 25 Hz for 30 sec in sterile 10 mM MgSO_4_. Dilutions of the plant homogenate (10 µL aliquots) were spotted on solid R2A media. The plates were incubated at 25°C for three days, and the number of colonies of each bacterial isolate was counted. Five technical replicate plates were analysed for each treatment and dilution and the experiment was carried out two times.

Colony forming unit (CFU) counts were Log_10_-transformed and LMs followed by ANOVA, and post-hoc analysis was performed with estimated marginal mean comparisons using the caret R package and emmeans R package. Electrolyte leakage values were normalised by arcsine transformation followed by pairwise comparison between bacterium-inoculated and mock-inoculated samples with Dunnett’s test using the DescTools R package (Signorell, 2023). Data were checked for normality and homoscedasticity using Shapiro‒Wilk and Levene’s test using the rstatix R package (Kassambara, 2023), respectively.

## Results

### Plant tissue, collection site and alpine Rosaceae plant are the main factors affecting the richness, diversity and taxonomic structure of endophytic bacterial communities

The analysis with high-throughput 16S rRNA amplicon sequencing of the endophytic bacterial communities associated with alpine Rosaceae plants (270 samples; Table S3) produced a total of 5,888 ASVs (13,074,650 filtered read counts; Table S4) with rarefaction curves that reached richness (observed ASVs) saturation (Fig. S2). In addition, a total of 152 ASVs were defined as the most abundant ASVs having a mean relative abundance of > 0.1% (Table S4).

Bacterial richness and alpha-diversity partially decreased from roots (average of 526.41 and 0.96) to leaves (average of 191.03 and 0.88) and to flowers (average of 179.24 and 0.86; Table S3). After splitting the data into two datasets (1 and 2), richness differed according to plant tissue (*F* = 290.76, *P* < 0.001) and collection site (*F* = 8.66, *P* < 0.001) in dataset 1 (*Alchemilla* sp. and *G. montanum* from six collection sites), as well as according to alpine Rosaceae plants (*F* = 10.00, *P* < 0.001), plant tissue (*F* = 83.33, *P* < 0.001) and collection site (*F* = 12.79, *P* < 0.001) in dataset 2 (*Alchemilla* sp., *D. octopetala* and *G. montanum* from two collection sites), according to LMs (Table S5). Alpha-diversity in dataset 1 and dataset 2 was influenced by plant tissue (*F* = 18.79, *P* < 0.001 and *F* = 5.04, *P* < 0.01, respectively), alpine Rosaceae plants (*F* = 6.18, *P* < 0.05 and *F* = 3.26, *P* < 0.05, respectively) and exposition (*F* = 6.71, *P* < 0.05 and *F* = 4.25, *P* < 0.05, respectively; Table S6). Collection site had also significant effect on alpha-diversity in dataset 1 (*F* = 3.19, *P* < 0.01). In particular, richness and alpha-diversity were higher in roots than in flowers and leaves according to estimated marginal mean comparisons in dataset 1 and dataset 2 (Fig. 1a and 1b; Tables S5 and S6). Among alpine Rosaceae plants, richness was higher in *Alchemilla* sp. compared to *G. montanum* and *D. octopetala* in dataset 2 (Table S5), while alpha-diversity was higher in *Alchemilla* sp. compared to *G. montanum* in dataset 1 and compared to *D. octopetala* in dataset 2 (Table S6). Among collection sites, richness and alpha-diversity were higher in the site D than in the other collection sites in dataset 1, and alpha-diversity was higher in samples from the North than in those from the South exposition in both datasets (Tables S5 and S6). Thus, plant tissue was also the main factor shaping richness and alpha-diversity in the conditional inference regression tree analysis (Fig. S3) and endophytic bacterial communities clustered according to plant tissue and alpine Rosaceae plant in the hierarchical clustering analysis (Fig. S4).

**Fig. 1.**
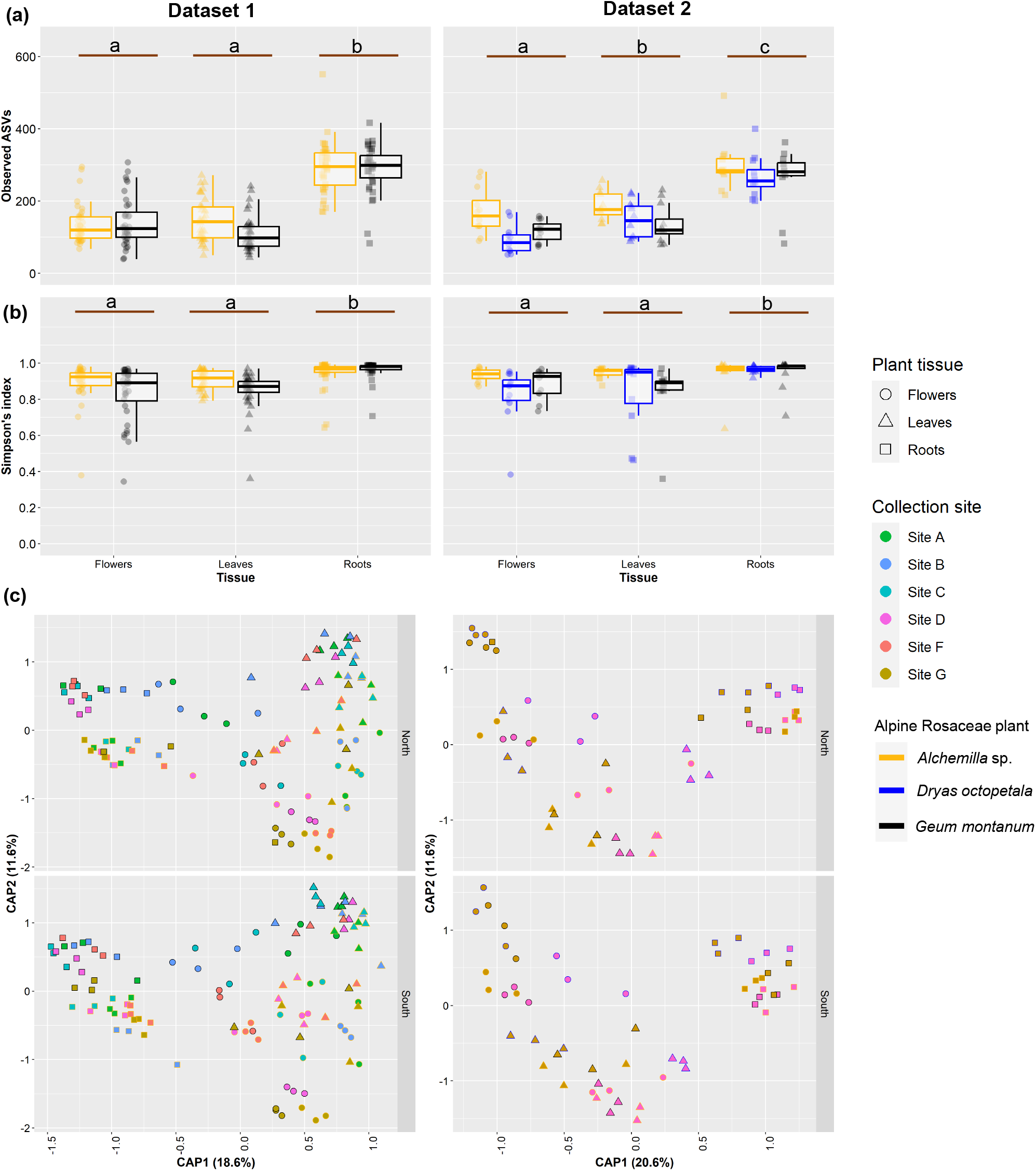
Diversity of endophytic bacterial communities associated with alpine Rosaceae plants. Observed amplicon sequence variants (ASVs) (richness) and Simpson’s index (alpha-diversity) are reported for the first dataset (a; dataset 1: *Alchemilla* sp. and *G. montanum* from six collection sites) and the second dataset (b; dataset 2: *Alchemilla* sp., *D. octopetala* and *G. montanum* from two collection sites) of samples. Lowercase letters indicate results of estimated marginal mean comparisons between tissues (Tables S5 and S6). Constrained analysis of principle (CAP) coordinates based on Bray–Curtis dissimilarity matrices on samples collected from flower, leaf and root tissues of alpine Rosaceae plants from different collection sites and exposition in dataset 1 and dataset 2 (c).

CAP ordinations based on Bray–Curtis dissimilarities demonstrated the differentiation of endophytic bacterial community structure according to plant tissue and alpine Rosaceae plant in both datasets (Fig. 1c). CAP permutation tests were used to assess the significance of constraints and showed that all factors (plant tissue, alpine Rosaceae plant, collection site and exposition) significantly affected the beta-diversity in both datasets (Table S7). In particular, the taxonomic structure of endophytic bacterial communities was mainly affected by plant tissue (*F* = 44.45; coefficient of determination (R^2^) = 0.21; *P* < 0.001 and *F* = 21.28, R^2^ = 0.18, *P* < 0.001), collection site (*F* = 6.26, R^2^ = 0.07, *P* < 0.001 and *F* = 16.18, R^2^ = 0.07, *P* < 0.001) and alpine Rosaceae plants (*F* = 15.49, R^2^ = 0.04, *P* < 0.001 and *F* = 7.52, R^2^ = 0.06, *P* < 0.001), with a minor contribution of exposition (*F* = 2.40, R^2^ = 0.01, *P* < 0.01 and *F* = 2.28, R^2^ = 0.07, *P* < 0.01) in dataset 1 and dataset 2, respectively, according to beta-diversity analysis carried out with PERMANOVA on Bray–Curtis dissimilarities (Table S8). A dispersion in the ASV data was found, particularly within plant tissues (*F* = 48.80, *P* = < 2.20e-16 and *F* = 5.12, *P* = 0.008 for dataset 1 and dataset 2, respectively), according to permutational multivariate analysis of dispersion (PERMDISP2; Table S8), suggesting that differences observed by PERMANOVA may be attributed to either centroid or dispersion. The mGLMs and model-based ordination on ASVs with an occupancy of 0.25 (131 and 161 ASVs in dataset 1 and dataset 2, respectively) supported the results obtained with PERMANOVA, and the variation in the taxonomic structure of endophytic bacterial communities was mainly explained by plant tissue (Dev = 4,193.22, *P* < 0.001 and Dev = 2,864.45, *P* < 0.001), followed by collection site (Dev = 2,898.54, *P* < 0.001 and Dev = 1,419.73, *P* < 0.001), alpine Rosaceae plant (Dev = 2,156.2, *P* < 0.001 and Dev = 1,935.43, *P* < 0.001) and exposition (Dev = 308.5, *P* < 0.001 and Dev = 279.5, *P* < 0.001) in dataset 1 and dataset 2, respectively (Fig. S5 and Table S9).

Within each plant tissue, endophytic bacterial communities differed according to alpine Rosaceae plant, collection site and exposition (Fig. S6 and Table S10). In particular, the HEEI values revealed that alpine Rosaceae plant influenced mainly root-associated communities (1.25 HEEI, *F* = 15.89, R^2^ = 0.15, *P* < 0.001 and 2.81 HEEI, *F* = 5.29, R^2^ = 0.20, *P* < 0.001), whereas collection site influenced mainly leaf-associated communities (0.37 HEEI, *F* = 4.6, R^2^ = 0.22, *P* < 0.001 and 1.48 HEEI, *F* = 7.39, R^2^ = 0.11, *P* < 0.001) and flower-associated communities (0.20 HEEI, *F* = 6.29, R^2^ = 0.26, *P* < 0.001 and 0.53 HEEI, *F* = 25.65, R^2^ = 0.30, *P* < 0.001) in dataset 1 and dataset 2, respectively (Table S10).

### Core endophytic bacterial taxa of alpine Rosaceae plants are selected by the host plant and the indicator taxa are enriched in flower and leaf tissues compared to roots

Endophytic bacterial communities associated with Rosaceae plants were dominated by Proteobacteria (80.5%), followed by Actinobacteria (9.61%), Bacteroidota (4.07%), Patescibacteria (2.84%), Chloroflexi (1.53%) and Deinococcota (0.53%) phyla in terms of relative abundance (Fig. S7a and S8a). Most taxa detected were shared among the three alpine Rosaceae plants, with differences in relative abundance among the three plant tissues. In particular, flower-associated communities were dominated by *Pseudomonadaceae* (46.84%), *Oxalobacteriaceae* (16.03%), *Burkholderiaceae* (8.09%) and *Rhizobiaceae* (5.96%); leaf-associated communities were dominated by *Enterobacteriaceae* (27.27%), *Oxalobacteriaceae* (22.58%), *Pseudomonadaceae* (16.03%) and *Burkholderiaceae* (4.47%); and root-associated communities were dominated by *Rhizobiaceae* (11.48%), *Pseudomonadaceae* (10.54%), *Oxalobacteriaceae* (7.14%) and *Microbacteriaceae* (6.8%) (Fig. S7b and S8b). At the genus level, flower-associated communities were dominated by *Pseudomonas* (46.8%), *Duganella* (7.93%), *Ralstonia* (6.27%), *Allorhizobium-Neorhizobium-Pararhizobium-Rhizobium* (5.78%) and *Buchnera* (3.1%); leaf-associated communities were dominated by *Escherichia-Shigella* (24.88%), *Pseudomonas* (16.03%), *Janthinobacterium* (12.22%), *Duganella* (3.65%), *Ralstonia* (3.5%) and *Sphingomonas* (2.27%); and root-associated communities were dominated by *Pseudomonas* (10.54%), *Allorhizobium-Neorhizobium-Pararhizobium-Rhizobium* (8.4%), *Flavobacterium* (3.7%), *Kineosporia* (3.37%), *Cryptosporangium* (2.37%) and *Galbitalea* (1.91%) (Fig. 2 and Fig. S8c).

**Fig. 2.**
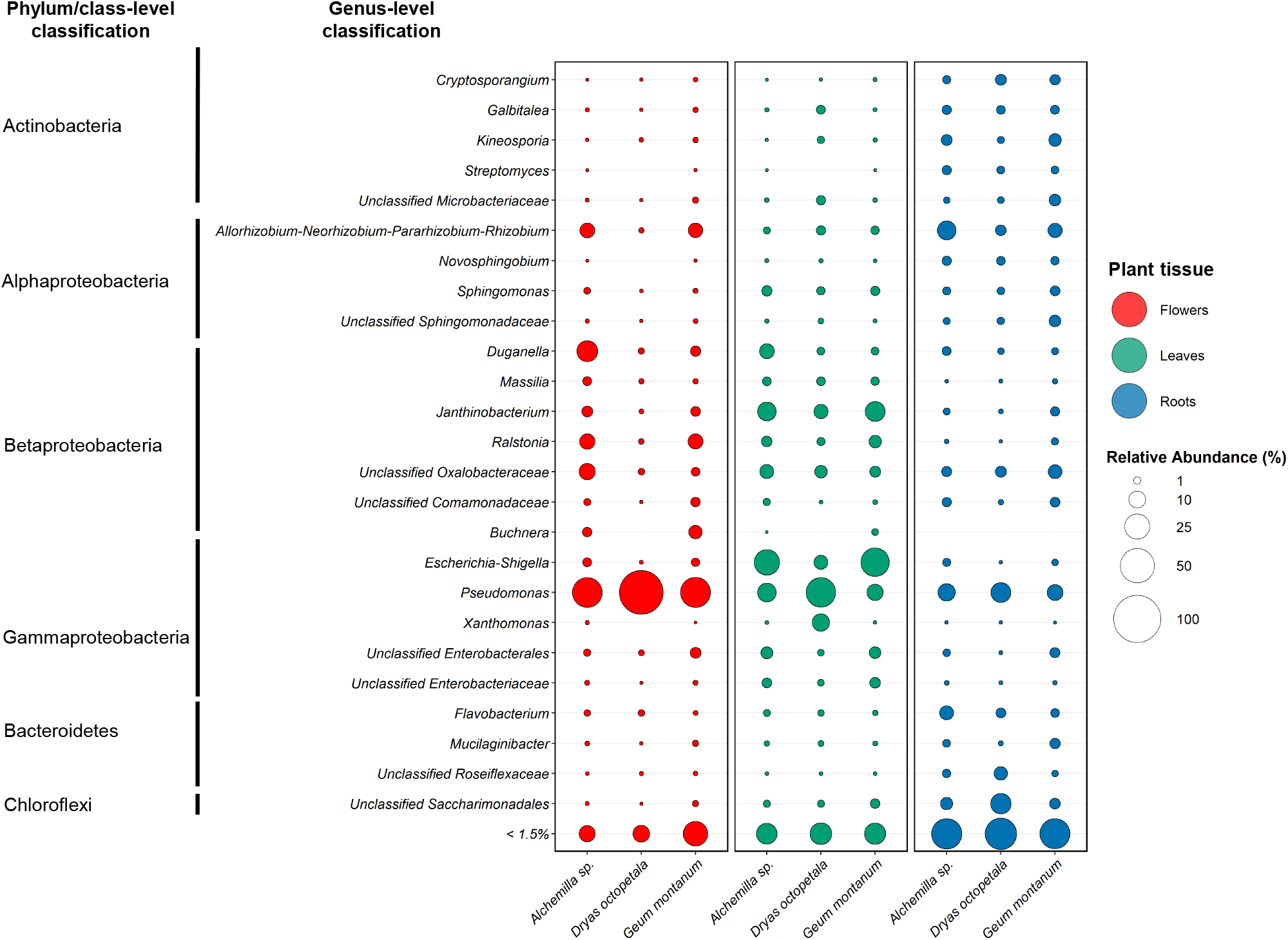
Taxonomic summary of the most abundant endophytic bacterial genera in alpine Rosaceae plants. Bubble plots represent the mean relative abundance of amplicon sequence variants (ASVs) aggregated to the genus level in each alpine Rosaceae plant and tissue across all samples, collection sites, and expositions. Taxa are coded with different colours according to the tissue.

According to the Sloan neutral model prediction (95% confidence intervals), 1,530 ASVs of the endophytic bacterial community fell above the prediction (higher occupancy than expected by their mean abundance; ASVs selected by the host or with good dispersal capabilities), 155 ASVs fell below the prediction (higher abundance than expected by their occupancy; ASVs not selected by the host or with scarce dispersal capabilities) and 4,203 ASVs followed the neutral process (Fig. 3a). In particular, 12 ASVs of the core endophytic bacterial taxa were above the neutral prediction, whereas 13 ASVs were below the predication. Core endophytic bacterial taxa of alpine Rosaceae plants were composed of 31 ASVs highly prevalent across all plant tissues, according to the abundance and occupancy distribution (Table S4). Although the core endophytic bacterial taxa represented a small fraction of the total community (0.53% of total ASVs), they accounted for 41% and 24% of the total relative abundance and beta-diversity, respectively, based on Bray–Curtis dissimilarities.

**Fig. 3.**
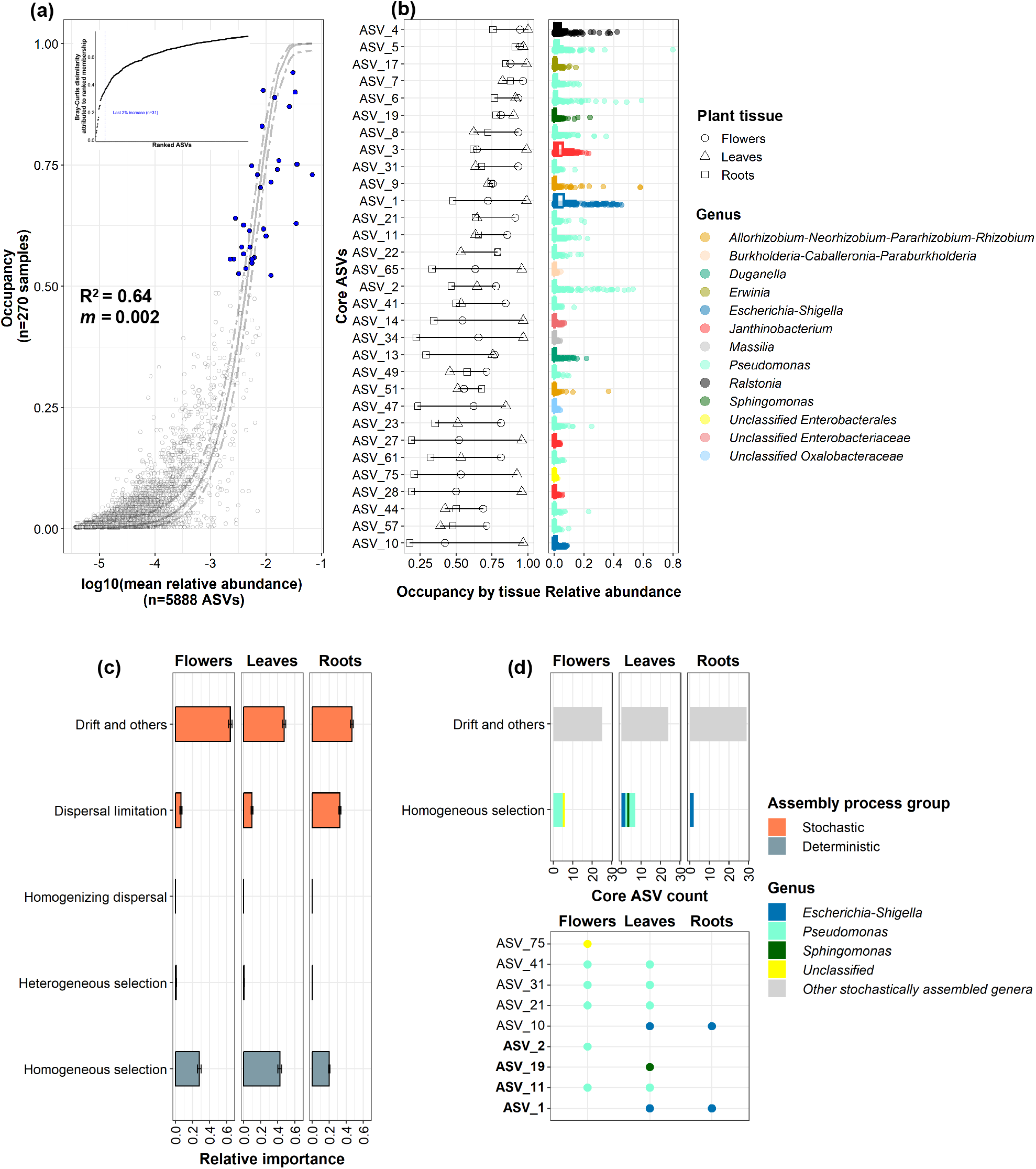
Abundance, occupancy and structure of core bacterial taxa in alpine Rosaceae plants. Abundance-occupancy distributions of amplicon sequence variants (ASVs) across flower, leaf and root tissue samples of *Alchemilla* sp., *Dryas octopetala* and *Geum montanum* (a). ASVs are represented by data points in grey and ASVs belonging to the core taxa are represented by blue dots. The solid line represents the fit of the Sloan neutral model, and the dashed line is the 95% confidence around the model prediction. R2 values (measurement of fit to neutral assembly process) and m-values (estimated migration rate) are indicated in the plot. The final 2% percent increase (represented by the blue dashed line) in beta-diversity based on Bray–Curtis dissimilarities due to the contribution of the prospective core set method used to identify core taxa is reported in the inset. The core taxa accounted for 24% of the total beta-diversity as shown in the inset. Occupancy, relative abundance and taxonomic annotation of ASVs of the core taxa are reported (b). The core taxa represented a small fraction (*n* = 31 ASVs) of the total community (*n* = 5,888 of total ASVs). The relative importance of stochastic and deterministic processes in the community assembly of endophytic bacterial communities was calculated in flower, leaf and root tissues (c). The ASV count and taxonomic annotation at the genus level are reported for the core taxa assembled through homogeneous selection (i.e. selection under homogeneous abiotic and biotic environmental conditions leading to more similar structures among communities) (d). Bold letters represent ASVs that were the top taxon within their respective phylogenetic bins.

The core endophytic bacterial taxa of alpine Rosaceae plants were composed of *Pseudomonas* (15 ASVs), *Janthinobacterium* (3 ASVs), *Escherichia-Shigella* (2 ASVs), *Allorhizobium-Neorhizobium-Pararhizobium-Rhizobium* (2 ASVs), *Burkholderia-Caballeronia-Paraburkholderia*, *Duganella*, *Erwinia*, *Massilia*, *Ralstonia* and *Sphingomonas* genera, with 1 ASV each, and unclassified taxa (3 ASVs; Fig. 3b). Moreover, the assembly of flower-, leaf- and root-associated communities was mainly driven by the stochastic processes of drift (relative importance of 0.64, 0.47 and 0.47, respectively) and dispersal limitation (relative importance of 0.07, 0.1 and 0.33, respectively) compared to the deterministic process of homogeneous selection (relative importance of 0.28, 0.42 and 0.2, respectively; Fig. 3c), and the strong stochastic contribution was confirmed by pNST analysis (Fig. S9). Within each plant tissue, the assembly of core endophytic bacterial taxa was mainly driven by the drift process (*n* = 27–29 ASVs) rather than homogeneous selection (*n* = 2–7 ASVs; Fig. 3d). Moreover, some core endophytic bacterial taxa (ASV 1, 2, 10, 11) resulted under homogeneous selection according to iCAMP model prediction (Fig. 3d), and they were the top taxa within their respective phylogenetic bins (relative abundance from 1 to 11% and relative importance from 0.02 to 23%; Table S11).

The indicator taxon analysis with random forest models revealed that 331 and 296 ASVs (indicator ASVs) contributed to the differences in community structure considering all factors (plant tissue, alpine Rosaceae plant, collection site and exposition) in dataset 1 and dataset 2, and they belonged to nine and eight genera, respectively (Table S12). Indicator ASVs belonging to the *Pseudomonas*, *Escherichia-Shigella*, *Duganella*, *Ralstonia* and *Janthinobacterium* genera were the most abundant indicator taxa (in terms of normalised read count), particularly in flower and leaf tissues rather than in root tissues (Fig. 4a). Differential abundance analysis on indicator ASVs confirmed that ASVs belonging to two (*Pseudomonas* and *Duganella*) and four (*Allorhizobium-Neorhizobium-Pararhizobium-Rhizobium*, *Kineosporia*, *Cryptosporangium* and *Galbitalea*) genera were enriched and depleted in flower and leaf tissues compared to root tissues, respectively (Fig. 4b and Table S13). Moreover, some indicator ASVs belonged to the core endophytic bacterial taxa, such as *Pseudomonas* (e.g. ASV 2, 5, 6, 8, 11, 21, 61), *Duganella* (ASV 13), *Erwinia* (ASV 17) and *Ralstonia* (ASV 4).

**Fig. 4.**
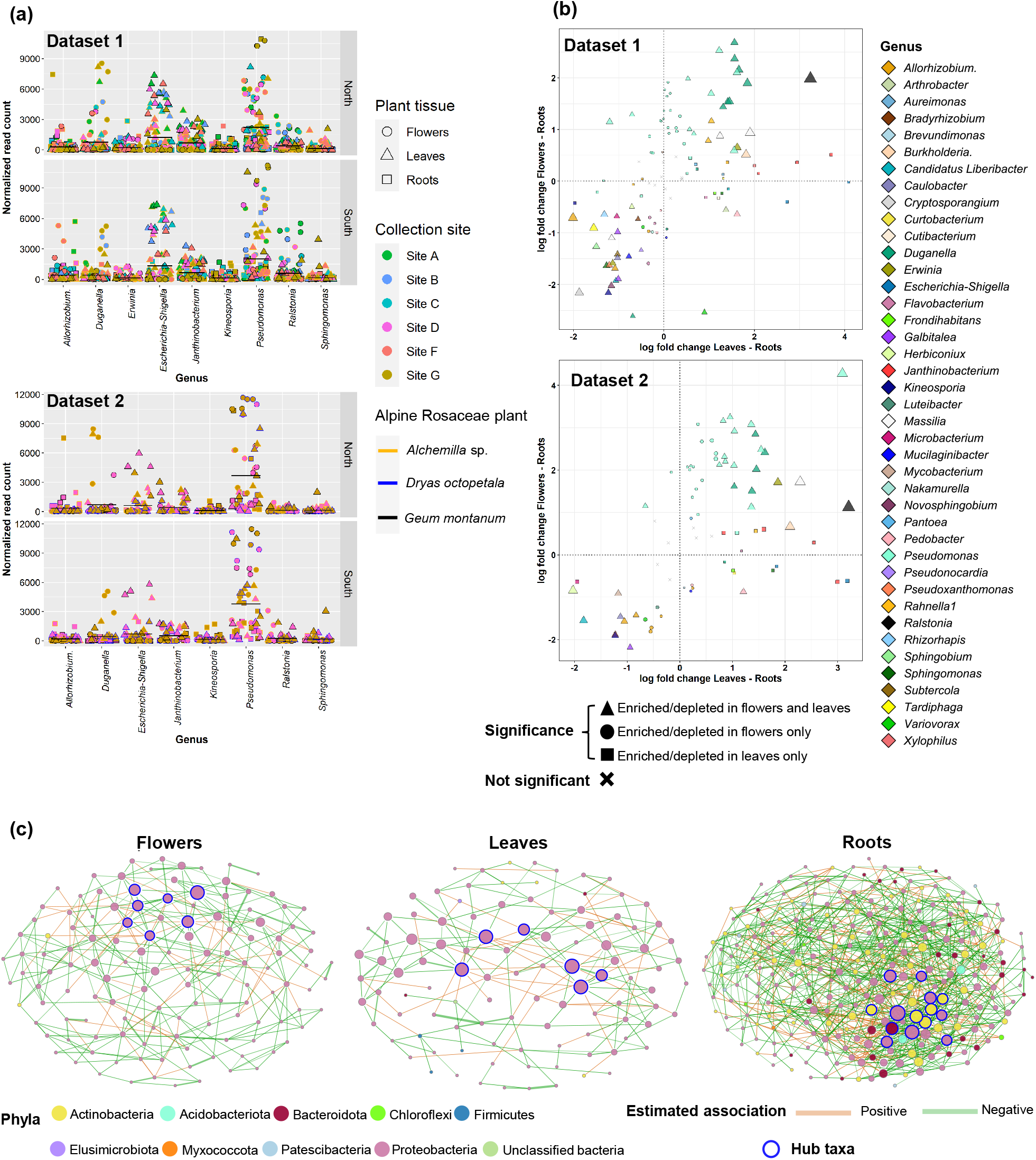
Identification of indicator and hub bacterial taxa enriched in different tissues of alpine Rosaceae plants. (a) A total of 331 and 296 amplicon sequence variants (ASVs) were identified as indicator ASVs by random forest models and were aggregated to 79 and 70 genera for the first dataset (*Alchemilla* sp. and *G. montanum* from six collection sites) and the second dataset (*Alchemilla* sp., *D. octopetala* and *G. montanum* from two collection sites), respectively. (b) Differentially abundant indicator ASVs are presented as log fold changes obtained from the log-linear model in flowers and leaves compared to roots. (c) Endophytic bacterial co-occurrence networks of Rosaceae plants. Each node corresponds to an ASV, and edges between nodes correspond to either positive (orange) or negative (green) correlations inferred from ASV abundance profiles. The thickness of each edge is proportional to the correlation coefficients of the connections. Node size reflects their eigenvector centrality, while nodes with blue outline colour represent hub taxa.

### Core endophytic bacterial taxa of alpine Rosaceae plants are hub taxa in flowers and leaves

A total of 128, 104 and 273 ASVs (nodes) were selected according to the relative abundance (> 0.001%) and occupancy (> 0.25) in flower, leaf and root tissues, respectively, and the co-occurrence network showed higher complexity and stronger connectivity in the roots than in the flowers and leaves (Fig. 4c; Table S14). In particular, higher number of hubs, higher average degree and lower average path distance were detected in roots (14 hubs, 7.28 average degree, 2.26 average path distance) than in flowers (7 hubs, 4.98 average degree, 2.36 average path distance) and leaves (6 hubs, 3.5 average degree, 2.82 average path distance). Comparative analysis of co-occurrence networks between similarity of most central nodes (i.e. set of ASVs with eigenvector centrality values greater than 95% of the empirical distribution of all eigenvector centralities in the network) showed that flower and leaf networks were more similar (from 0.098 to 0.61 Jaccard index) compared to flower and root networks (from 0.0 to 0.194 Jaccard index) or leaf and root networks (from 0.0 to 0.267 Jaccard index) (Fig. S10 and Table S14). Some ASVs of flower (ASV 1 and 11) and leaf (ASV 2, 4, 7, 10) hub taxa were also classified as core endophytic bacterial taxa, while root hub taxa comprised non-core taxa, such as *Sphingomonadaceae* (ASV 560, 280), *Microbacteriaceae* (ASV 200, 206) and *Sphingobacteriaceae* (ASV 314) (Fig. 4c and Table S14).

### Culturable bacterial endophytes belonging to the core endophytic bacterial taxa mitigate freezing stress in strawberry seedlings

To test the functional role of bacterial endophytes of the alpine Rosaceae plants belonging to the core taxa, a large number of culturable psychrotolerant bacteria (*n* = 685 isolates) were isolated from the above-described plant samples under the selected isolation conditions. The isolates belong to four phyla and 65 genera and were isolated from *Alchemilla* sp. (*n* = 296), *D. octopetala* (*n* = 132) and *G. montanum* (*n* = 257) (Fig. 5a and Table S15). The collection was mainly composed of genera belonging to Proteobacteria, such as *Pseudomonas* (*n* = 314), *Sphingomonas* (*n* = 93), *Rhizobium* (*n* = 35), *Rahnella* (*n* = 27), *Erwinia* (*n* = 26) and *Duganella* (*n* = 16), and Bacteroidota, such as *Pedobacter* (*n* = 27), *Mucilaginibacter* (*n* = 16), *Flavobacterium* (*n* = 12), Actinobacteria (*n* = 37) and Firmicutes (*n* = 7). The density of culturable psychrotolerant bacterial endophytes varied according to plant tissue, collection site and alpine Rosaceae plant, and it was higher in root and flower tissues than in leaf tissues (Fig. S11 and Table S16). The results from the sequence search of the 16S rRNA

**Fig. 5.**
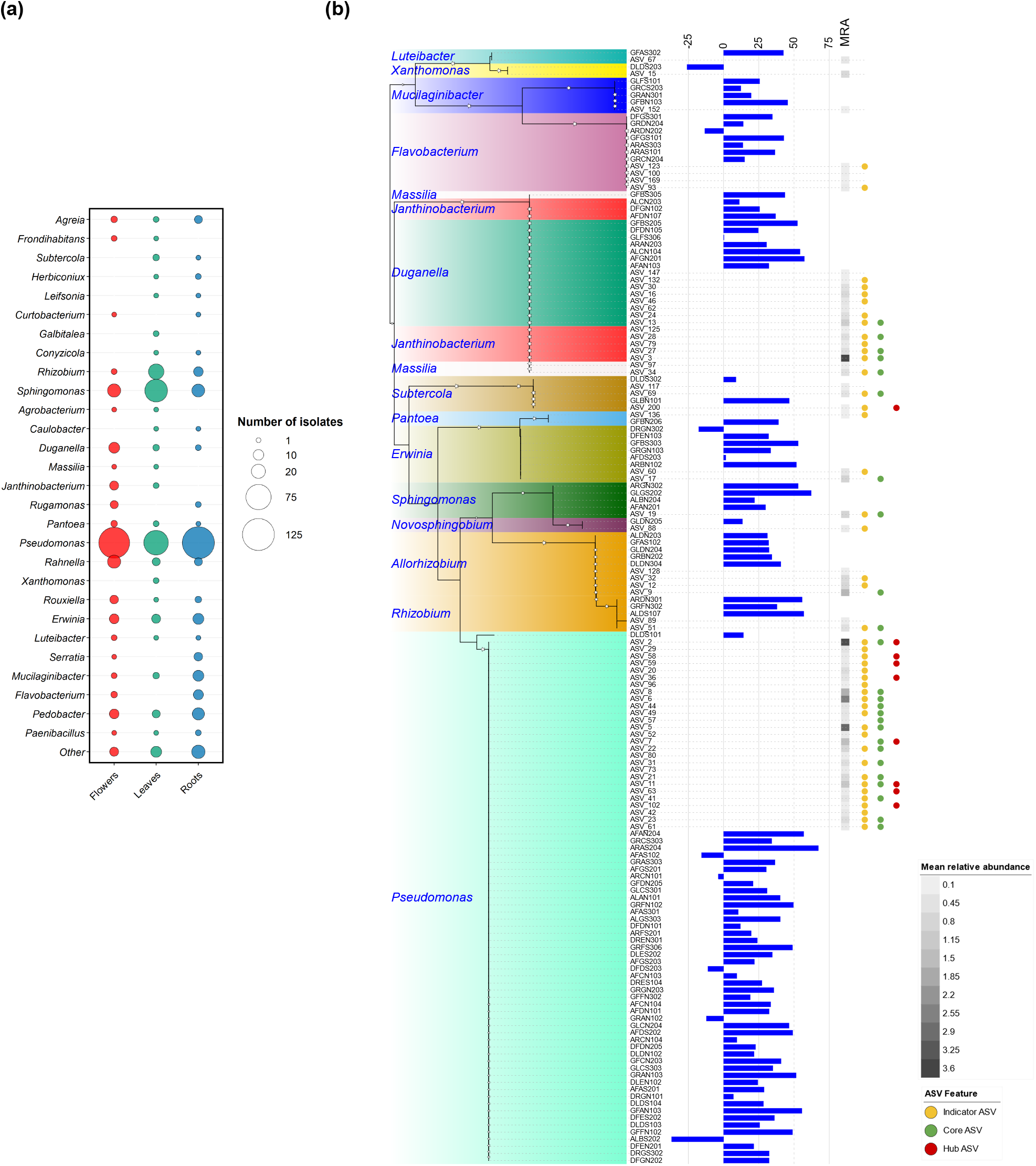
List of the psychrotolerant bacterial endophytes isolated from the selected alpine Rosaceae plants and their freezing protection ability. (a) Taxonomic summary of the culture collection of psychrotolerant bacterial endophytes (685 isolates) isolated from flowers, leaves and roots of *Alchemilla* sp., *Dryas octopetala* and *Geum montanum.* (b) Phylogenetic tree of representative psychrotolerant bacterial endophytes (94 isolates) and the most abundant amplicon sequence variants (63 ASVs with mean relative abundance > 0.1%). For each isolate, the blue bar represents the reduction in electrolyte leakage (%) obtained in bacterium-inoculated compared to mock-inoculated strawberry plants after freezing stress. For each ASV, grey square indicates the mean relative abundance of ASVs across all samples according to the scale legend. Coloured circles indicate indicator taxa (yellow), core taxa (green) and hub taxa (red).

Sanger sequences of culturable psychrotolerant bacterial endophytes against all recovered ASVs (35,300 ASVs before any filtering) revealed 94 isolates with more than 99% nucleotide identity with 63 most abundant ASVs (mean relative abundance > 0.1%) and they were defined as representative psychrotolerant bacterial endophytes (Fig. 5b; Table S17). The majority (91.5%) of representative psychrotolerant bacterial endophytes were able to reduce electrolyte leakage in strawberry seedlings exposed to freezing stress and the best performing isolates (reduction in electrolyte leakage > 50%) belonged to the core endophytic bacterial taxa of alpine Rosaceae plants, such as *Duganella*, *Erwinia*, *Pseudomonas, Rhizobium* and *Sphingomonas* (Fig. 5b). In the validation tests, *Duganella* sp. GFBS205, *Duganella* sp. ALCN104, *Erwinia* sp. GFBS303, *Pseudomonas* sp. AFDS202 and *Rhizobium* sp. ALDS107 confirmed the reduction in electrolyte leakage in strawberry seedlings exposed to freezing stress (Fig. 6). These bacterial isolates were able to colonize strawberry seedlings (Fig. 6) and to survive at −6°C on treated plants (data not shown).

**Fig 6.**
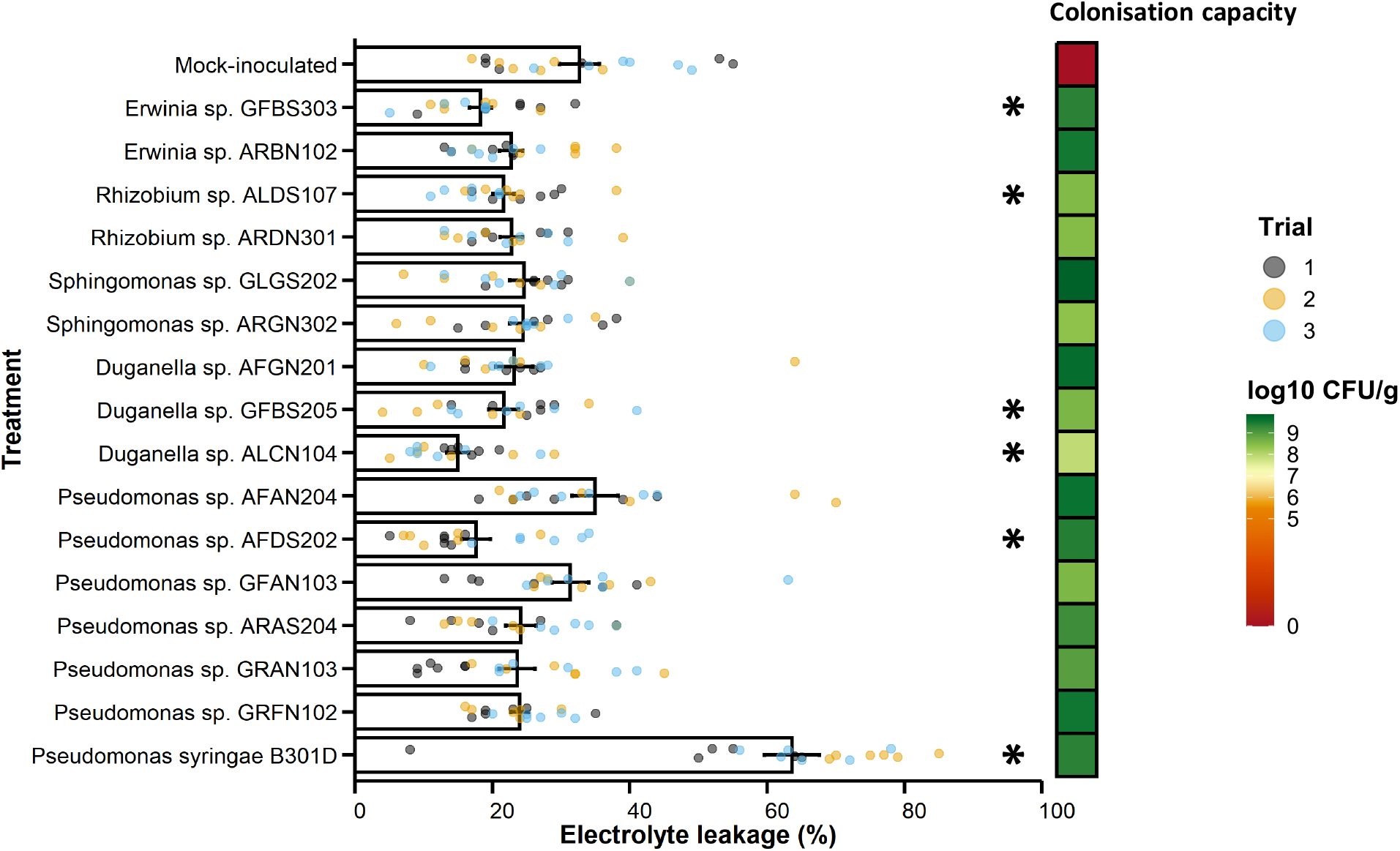
Freezing protection ability of the selected psychrotolerant bacterial endophytes isolated from the alpine Rosaceae plants. Electrolyte leakage was used to assess freezing damage in strawberry seedlings treated with MgSO_4_ (mock-inoculated) or inoculated with selected psychrotolerant bacterial endophytes of alpine Rosaceae. Significant differences between bacterium-inoculated and mock-inoculated samples are indicated by asterisks according to Dunnett’s test (*P* < 0.05). The colonisation capacity of the isolates was checked by dilution-plating and represented as a heatmap. Data from three independent experiments (represented by different colours) with six replicates (seedlings) each are reported. *Pseudomonas syringae* B301D was used as a control for its ability to increase freezing damage.

## Discussion

### Endophytic bacterial communities of alpine Rosaceae plants are affected by the plant tissue, collection site and host plant

Endophytic bacterial communities associated with alpine Rosaceae plants were dominated by *Pseudomonadaceae* and *Oxalobacteriaceae* families across all three plant tissues (flower, leaf and root), while enriched families were found in each plant tissue, such as *Burkholderiaceae* in flowers, *Enterobacteriaceae* in leaves, and *Rhizobiaceae* and *Microbacteriaceae* in roots. Similarly, several of these bacterial families are among the dominant endophytic taxa of other alpine plants, arctic plants (e.g. *Oxyria digyna* and *Saxifraga oppositifolia*) (Kumar *et al*., 2017; Brigham *et al*., 2023) and many agronomic plants (e.g. canola, rapeseed, strawberry, tobacco and tomato and wheat) (Dong *et al*., 2019; Rathore *et al*., 2019; Chen *et al*. 2020; Zhang *et al*., 2020a; Elvia *et al*., 2021), suggesting that such bacterial taxa are associated with plants.

Plant-associated bacterial endophytes originate from soils, rainfall and aboveground plant niches, such as the phyllosphere (leaf surface) and anthosphere (the external environment of flowers), via wind- or insect-mediated transmission and, following their entrance, bacteria can spread systemically via the xylem to aboveground tissues (Compant *et al*., 2019). Bacterial richness and alpha-diversity partially decreased from roots to leaves and flowers of alpine Rosaceae plants and only a small fraction of ASVs (less than 7%) was shared between the aboveground tissues and root tissues, suggesting a contribution of tissue-specific factors in endophytic bacterial community shaping, such as physical and chemical barriers, root exudates and flower and leaf metabolites. Thus, the plant tissue explained the greatest amount of variation (approximately 18–21% explained variance) in the community structure of alpine Rosaceae plants. Likewise, plant tissue showed a strong selective force in shaping plant-associated microbial communities in Antarctica and Arctic regions (Given *et al*., 2020; Zhang *et al*., 2020b), indicating that plant tissue could be a major factor affecting bacterial community structure in cold environments. In particular, ASVs belonging to the genera *Pseudomonas* (*Pseudomonadaceae*), *Escherichia-Shigella* (*Enterobacteriaceae*), *Duganella* and *Janthinobacterium* (*Oxalobacteriaceae*) and *Ralstonia* (*Burkholderiaceae*) were indicator taxa with higher abundance in aboveground tissues (flower and/or leaf tissues) compared to root tissues of alpine Rosaceae plants, indicating the possible recruitment of well-adapted bacterial taxa, such as *Escherichia-Shigella, Pseudomonas* and *Ralstonia* previously found in aboveground tissues of strawberry and apple (Wei & Ashman 2018; Zhang *et al*., 2020a; Longa *et al*., 2022). The effect of the plant tissue on the structure of endophytic bacterial communities indicated niche-partitioning (i.e. separation among bacterial taxa in the tissue), possibly due to the variation in chemical and physical characteristics of each tissue (Compant *et al*., 2021).

In addition to the plant tissue, collection site (7% explained variance), alpine Rosaceae plants (5% explained variance) and exposition (less than 1% explained variance) affected the community structure of endophytic bacterial communities, which could be related to the different soil and host properties and climatic conditions. Likewise, previous studies reported the influence of environmental-related factors (e.g. soil fraction, soil texture, soil pH and snow depth) and plant-related factors (e.g. as plant species and plant neighbourhood in terms of plant composition and density) on the structure of plant-associated bacterial communities in alpine regions (Massaccesi *et al*., 2015; Řeháková *et al*., 2015; Chang *et al*., 2018; Wassermann *et al*., 2019; Brigham *et al*., 2023). For example, Brigham *et al*. (2023) showed that plant species are the major driver of endophytic bacterial communities, indicating that multiple factors contribute to defining the taxonomic structure of endophytic bacterial communities of alpine Rosaceae plants. In particular, bacterial community assembly is governed by two complementary mechanisms of stochastic and deterministic processes (Zhou & Ning 2017). Stochastic processes may include drift (random changes in population size due to birth and death events), dispersal (movement of microorganisms between different locations) and diversification (evolutionary process that generates genetic variation), while deterministic processes may include selection (biotic and abiotic interactions resulting in fitness differences) (Zhou & Ning 2017). In alpine Rosaceae plants, the contribution in the assembly of endophytic bacterial communities was higher for stochastic processes of drift and dispersal compared to deterministic processes of homogeneous selection (i.e. tissue selection). Likewise, the main role of stochastic processes in shaping the bacterial community structure was previously found in alpine plants (e.g. *Elymus nutans*, *Festuca sinensis*, *Kobresia pygmaea* and *Kobresia humilis*) (Liu *et al*., 2021; Wei *et al*., 2022) and it is possibly associated with competitive exclusion, historical contingency (i.e. result of a disturbance effect, such as recurrent frosts) and adaptability of bacterial endophytes to variation in environmental conditions, as suggested for other plant hosts (Cheng *et al*., 2023; Cregger *et al*., 2018; Hough *et al*., 2020; Xie *et al*., 2021; Zhu *et al*., 2021; Bell *et al*., 2022; Taniguchi *et al*., 2023). However, the assembly of plant-associated bacterial communities can shift between stochastic and deterministic processes depending on plant genotype, age and growth stage (Dove *et al*., 2021; Bell *et al*., 2022; Wang *et al*., 2023), indicating the possible contribution of both processes.

### Core endophytic bacterial taxa of alpine Rosaceae plants are hub taxa

The core endophytic bacterial taxa of alpine Rosaceae plants were composed of 31 ASVs belonging to the genera *Allorhizobium-Neorhizobium-Pararhizobium-Rhizobium*, *Burkholderia-Caballeronia-Paraburkholderia*, *Duganella, Erwinia, Escherichia-Shigella*, *Janthinobacterium*, *Massilia, Pseudomonas*, *Ralstonia* and *Sphingomonas*. These taxa were previously found as a part of the core microbiome in seeds, leaves and roots of alpine plants in the Andes, Qinghai Tibet Plateau (China), Rocky Mountains (USA) and European Alps (Ruiz-Perez *et al*., 2016; Kumar *et al*., 2017; Wassermann *et al*., 2019; Liang *et al*., 2021; Brigham *et al*., 2023), suggesting the presence of ubiquitous core taxa associated with alpine plants across divergent plant species, plant tissues and environmental conditions. Although most of the core taxa were assembled through the drift process, several ASVs belonging to *Pseudomonas* were deterministically assembled by homogeneous selection in the aboveground tissues, suggesting their selective recruitment by alpine Rosaceae plants.

Hub taxa can be key drivers in the microbial community structure and function of a host plant, particularly if they are part of the core microbiome and are consistently present in an environment (Agler *et al*., 2016; Banerjee *et al*., 2018). In alpine Rosaceae plants, hub taxa were found as part of the core taxa in aboveground tissues (flowers and leaves), but not in root tissues, and they belonged to the genera *Escherichia-Shigella*, *Pseudomonas* and *Ralstonia*. Likewise, the genera *Escherichia-Shigella*, *Pseudomonas* and *Ralstonia* were previously found as hub taxa of plant- and soil-associated microbial communities in agricultural lands and extreme environments (Li *et al*., 2022; L. Zhang *et al*., 2022a, X. Zhang *et al*., 2022b; Zheng *et al*., 2022; Zhu *et al*., 2023), suggesting that members of these genera may be adapted to a wide range of host plants and environmental conditions with a key contribution to defining microbial community structure and function. In particular, the strong interaction of hub taxa with other bacterial taxa in the co-occurrence networks of alpine Rosaceae plants revealed increased (e.g. nodes, hubs and average degree) and decreased (e.g. average path distance) topological properties in roots compared to flowers and leaves, suggesting strong complexity, connectivity and bacterial interactions in roots, as previously reported for various plants (e.g. strawberry) (Xiong *et al*., 2021; Zhang *et al*., 2022b; Hassani *et al*., 2023).

### Culturable psychrotolerant bacterial endophytes contribute to plant freezing stress tolerance

Culturable psychrotolerant bacterial endophytes represent a selection of endophytic bacterial communities associated with alpine Rosaceae plants and they succeed in mitigating freezing stress in strawberry seedlings. In particular, *Erwinia* (*Erwiniaceae*), *Duganella* (*Oxalobacteriaceae*), *Pseudomonas* (*Pseudomonadaceae*) and *Rhizobium* (*Rhizobiaceae*) isolates reduced electrolyte leakage of strawberry seedlings exposed to freezing stress and belonged to core endophytic bacterial taxa according to bioinformatics analysis. The functional importance of core bacterial taxa was previously shown for maize growth (Zhang *et al*., 2022a) and banana health (Hong *et al*., 2023), suggesting that core bacterial taxa are selected by the host plant to provide beneficial effects on plant growth and survival under stress conditions. Likewise, previous works showed that *Pseudomonadaceae* and *Oxalobacteraceae* families were enriched under cold stress in cold-sensitive (e.g. sorghum, taxus, lettuce and maize) and cold-adapted (e.g. *Valerianella locusta*) plants (Yu *et al*., 2019; Erlandson *et al*., 2021; Persyn *et al*., 2022; Garcia Mendez *et al*., 2023). Moreover, bacterial isolates belonging to *Pseudomonadaceae, Oxalobacteraceae and Rhizobiaceae* promoted plant growth under cold stress and/or reduced cold stress-related damage in several plants (Bertrand *et al*., 2007; Tiryaki *et al*., 2019; Acuña-Rodríguez *et al*., 2020; Beirinckx *et al*., 2020; D’Amours *et al*., 2022; Garcia Mendez *et al*., 2023). For example, inoculation with rhizobia (*Sinorhizobium meliloti*) led to a high proportion of undamaged nodules after freezing stress in alfalfa (D’Amours *et al*., 2022) and inoculation with *Pseudomonas* spp. (lacking the *ina* gene responsible for ice nucleation) improved freezing stress tolerance in Rosaceae plants (Hirano & Upper 2000; Tiryaki *et al*., 2019), corroborating that some plant-associated bacterial taxa contribute to cold stress mitigation.

Some *Pseudomonas*-based biostimulants for the mitigation of frost damage are already available on the market, mostly applied to vegetative plant tissues (e.g. leaves) to control *ina*-producing bacterial populations (Fadiji *et al*., 2022). Our discovery of the potential of *Pseudomonas* and non-*Pseudomonas* in freezing stress mitigation expands on this by offering isolates acting through mechanisms other than controlling *ina*-producing bacteria. The next step would be evaluating the effect and mode of action of the most promising isolates on vulnerable plant tissues, such as flowers, to further develop a cost-effective strategy to mitigate spring frost on fruit tree crops.

## Conclusion

In agreement with previous studies on other species, we found that the selected Alpine Rosaceae plants of this study are colonised by complex endophytic bacterial communities whose taxonomic structures are affected by plant tissue, collection site and host plant. These communities are assembled through stochastic processes of drift and deterministic processes of homogeneous selection and the identified core, hub and indicator endophytic bacterial taxa could play important functions in alpine Rosaceae plants. In particular, we found that culturable psychrotolerant bacterial endophytes isolated from these alpine Rosaceae plants can mitigate freezing stress in strawberry plants, suggesting that some members of endophytic bacterial communities can contribute to cold stress mitigation in these plants. The bacterial isolates that provided efficient freezing mitigation belonged to *Duganella, Erwinia, Pseudomonas* and *Rhizobium*, indicating that functional characterisation of core and hub endophytic bacterial taxa is the starting point for further manipulation of the plant microbiome under field conditions through a stable synthetic community or for the development of efficient microbial biostimulants for cold stress mitigation in agriculture. Future studies should aim to understand the factors affecting the functionality of core microbiota as well as the mode of action, effective target site and field performance of the most promising psychrotolerant bacterial endophytes.

## Acknowledgements

This work was funded by the European Union’s Horizon 2020 Research and Innovation Program Under the Marie Skłodowska Curie grant agreement number 101021787 (project FreezingBioprotector). We thank Melissa Alussi and Carmela Sicher from Fondazione Edmund Mach for providing help in the functional analysis of bacteria.

## Author contribution

MM and MP conceptualized and designed the experiment and coordinated all research activities. MM and MP collected the samples and performed DNA extraction. MM isolated culturable bacteria and prepared the samples for amplicon sequencing. MM characterised the bacterial isolates. MM, MP, LA performed the biostatistical analysis. MM and LA analysed the amplicon sequencing data. MM, MP, IP, and LA contributed to data interpretation and manuscript writing. All the authors revised and approved the final manuscript.

## Data availability

Amplicon sequencing data are available from the NCBI Sequence Read Archive under the BioProject number PRJNA872124. Sanger sequencing data are available under the NCBI GenBank accession numbers OR809314–OR809998 (accession number for each bacterial isolate are reported in Table S15).

## Ethical approval

All the field sampling of Rosaceae plants were carried out in agreement with institutional, national, and international guidelines and legislation.

## Supplementary figures

**Fig S1.**
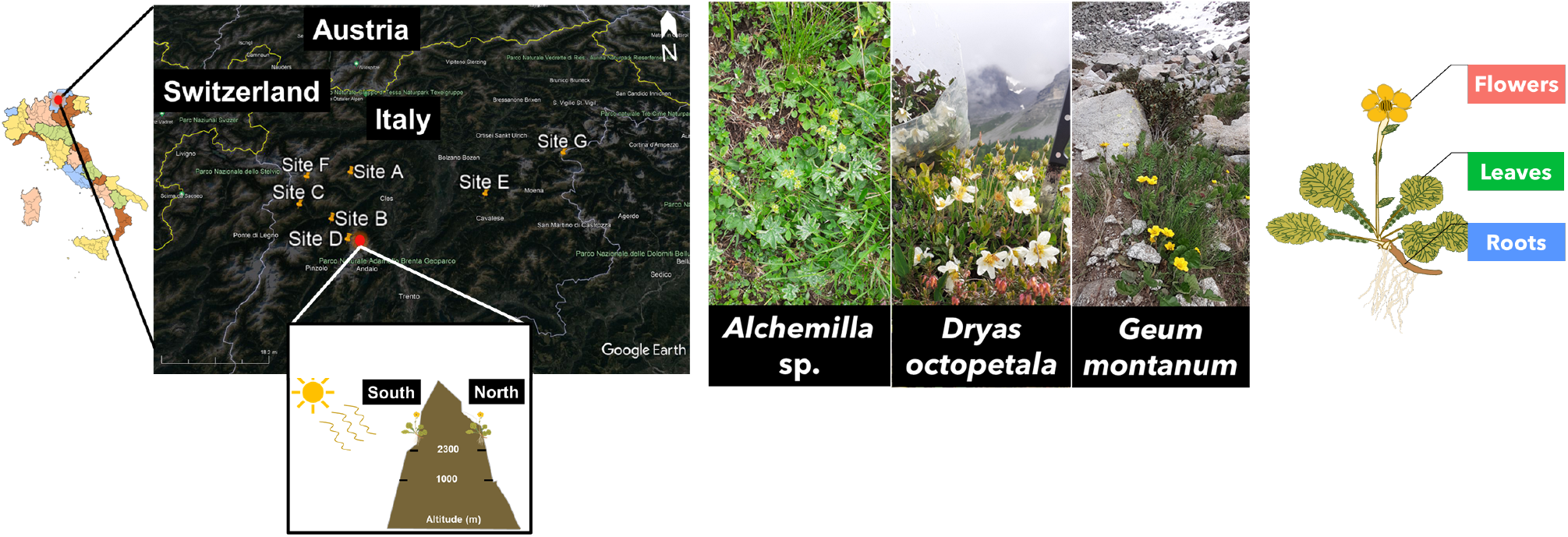
Experimental design of the study. A total of 270 samples were analysed: 108 from *Alchemilla* sp., 54 from *Dryas octopetala* and 108 from *Geum montanum* from three tissues, from seven sites and two expositions.

**Fig S2.**
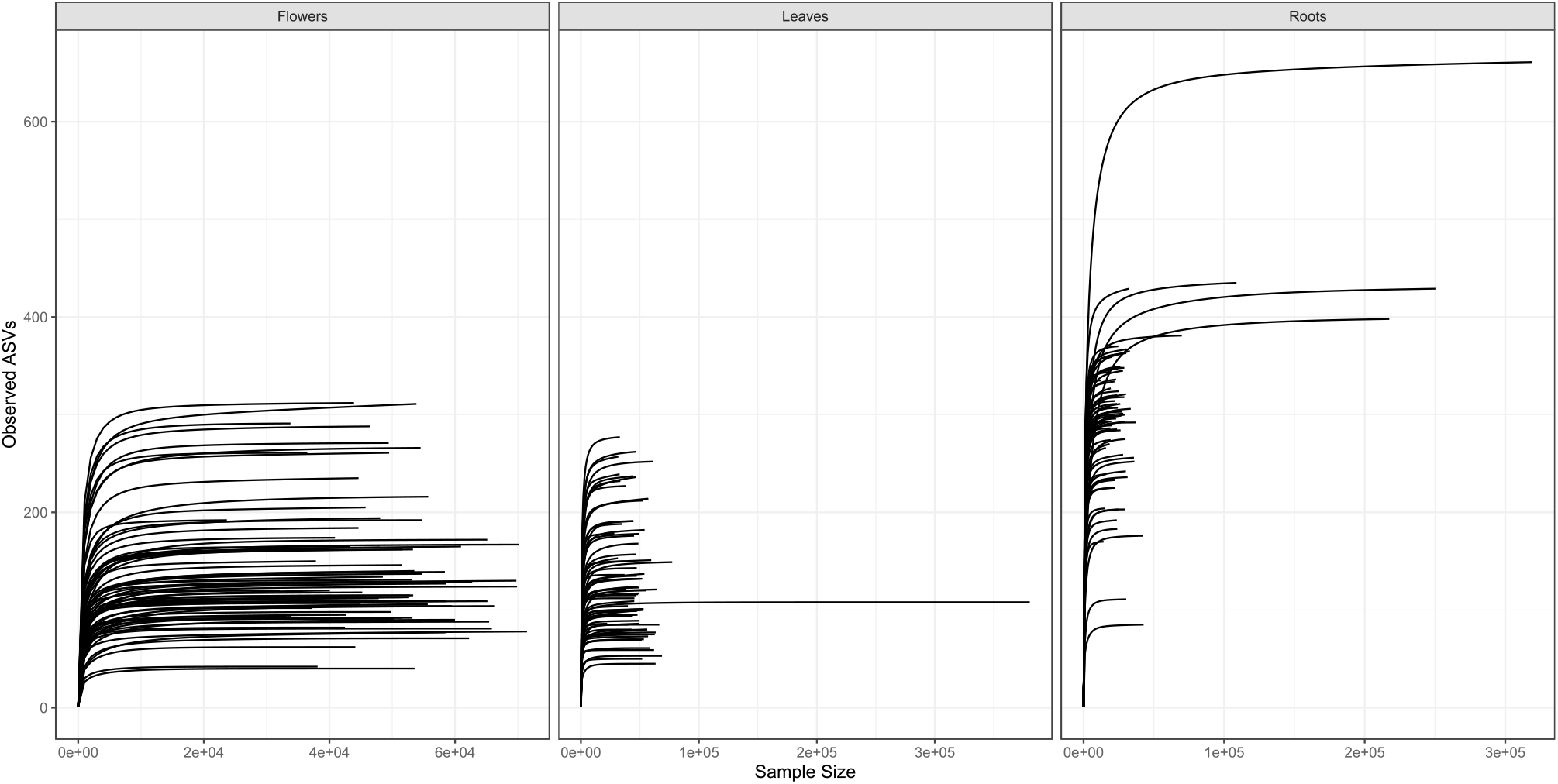
Rarefaction curves of endophytic bacterial communities associated with alpine Rosaceae plants. Curves were generated using the rarecurve function with 100 step size from the vegan R package. A total 270 samples; 108 from *Alchemilla* sp., 54 from *Dryas octopetala* and 108 from *Geum montanum* from flower, leaf and root tissues from seven collection sites and two expositions were analysed.

**Fig S3.**
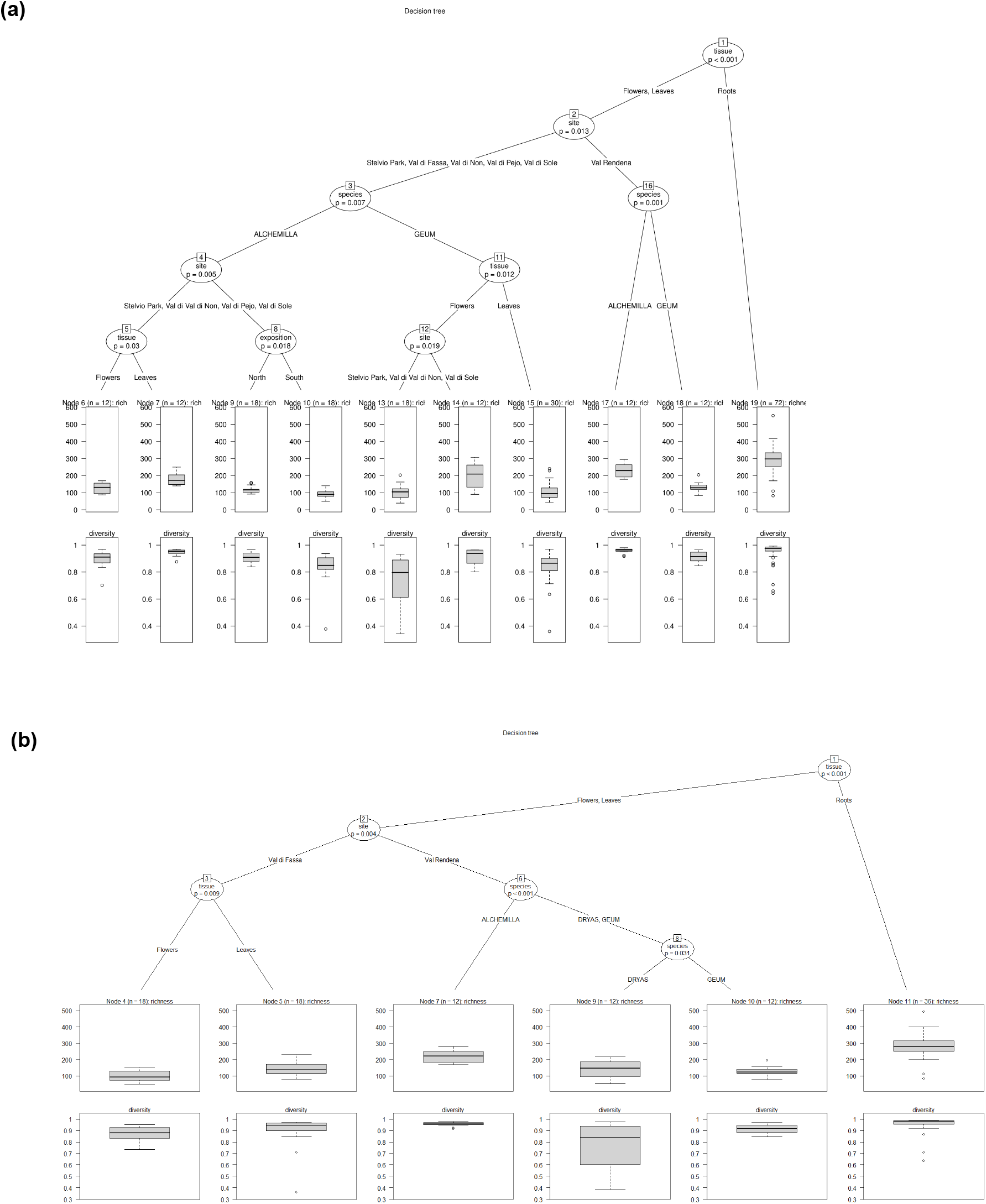
Conditional inference regression tree of bacterial alpha-diversity. Conditional inference regression trees were fitted onto the richness (observed ASVs) and diversity (Simpson’s index) values from the first dataset (a; *Alchemilla* sp. and *G. montanum* from six sites) and the second dataset (b; *Alchemilla* sp., *D. octopetala* and *G. montanum* from two sites) of samples to test the global null hypothesis of independence for covariate variables (i.e. tissue, species, site and exposition), using the ctree function from partykit v1.2 R package. Each node (circles) represents the variable that was split and gives p-values (determined using Bonferroni corrected p-values from permutation tests) associated with the split. Factors that were split for each variable are shown below each circle. Terminal nodes at the bottom of each tree include the mean alpha-diversity metrics of richness (top boxes) and diversity (bottom boxes) grouped within the splitting criteria.

**Fig S4.**
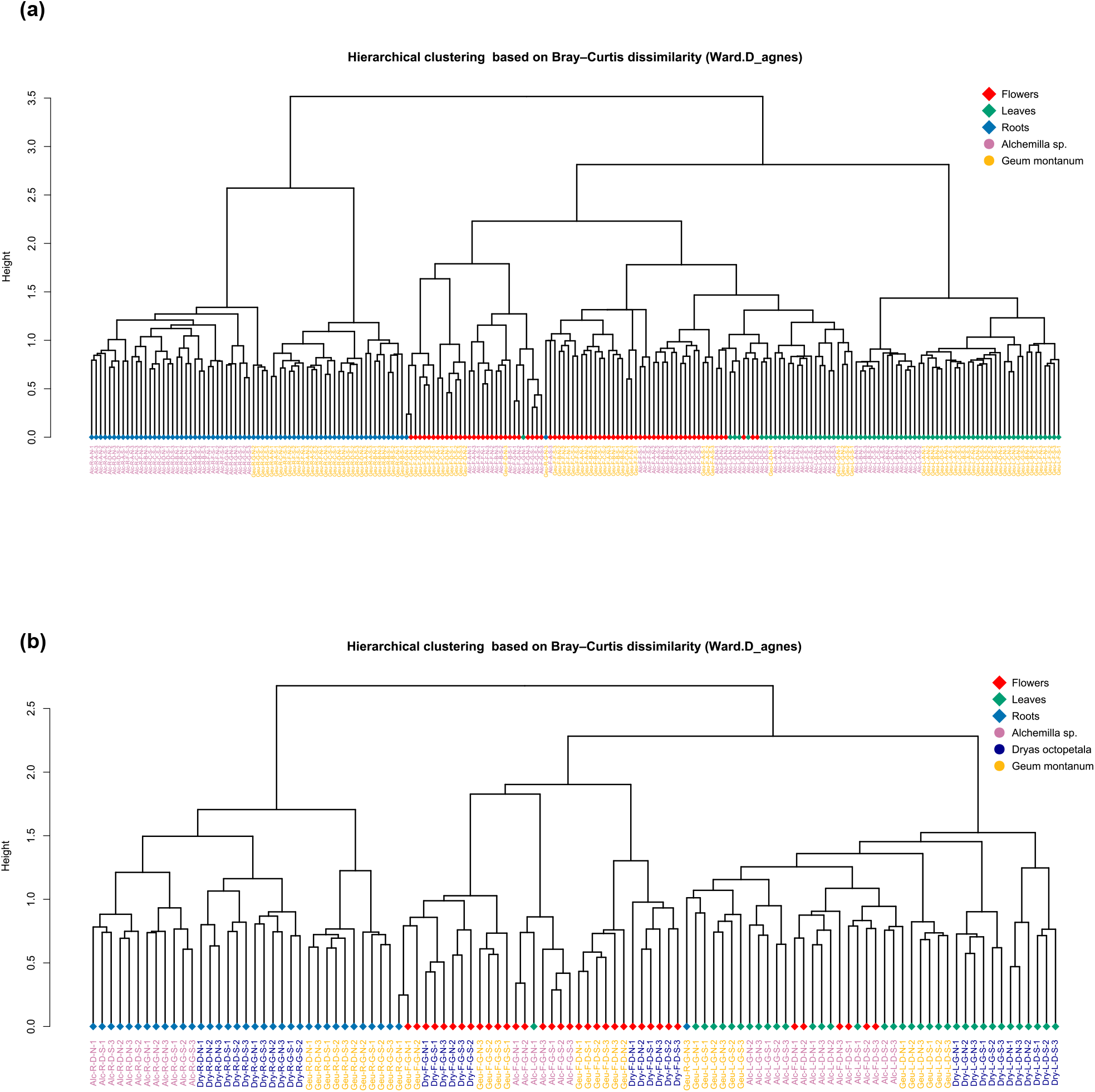
Hierarchical clustering of alpine Rosaceae plants samples. Hierarchical clustering of the first dataset (a; *Alchemilla* sp. and *G. montanum* from six collection sites) and the second dataset (b; *Alchemilla* sp., *D. octopetala* and *G. montanum* from two collection sites) of samples was based on Bray–Curtis dissimilarity matrices obtained from rarefied and Wisconsin-double standardised count data. Hierarchical clusters were constructed using agnes function with Ward.D method (identified as the strongest clustering structure by agglomerative coefficient test).

**Fig S5.**
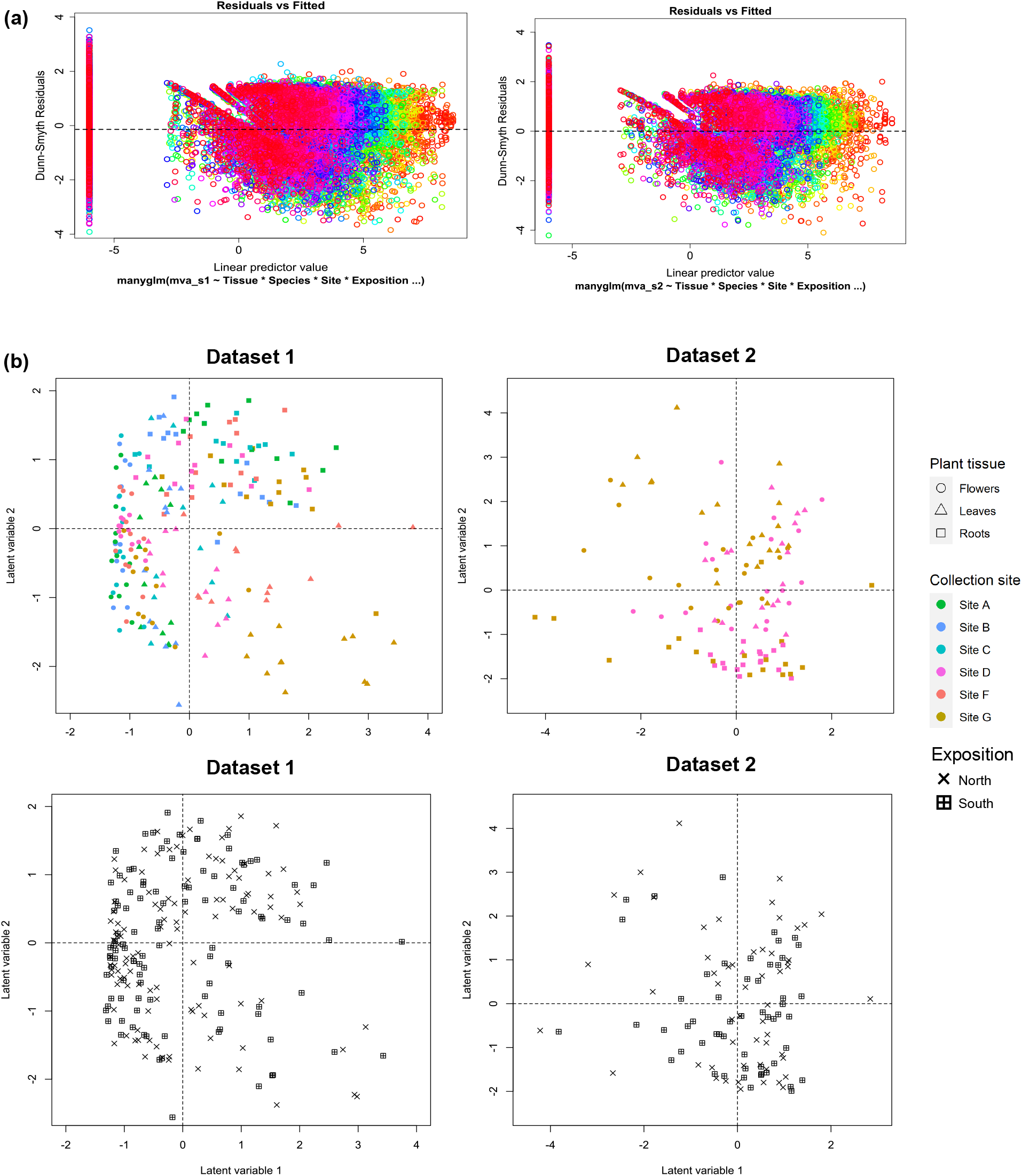
Multivariate generalized linear model (GLM) analysis on endophytic bacterial community structure of alpine Rosaceae plants. The residual plot shows the negative binomial distribution of the data and the ordination biplot displays the endophytic bacterial community structure of Rosaceae plants (a). A model-based ordination with Gaussian copulas to visualize the GLM results (obtained through mvabund package) were computed using the ordiplot function from the gllvm R package (b). The first dataset (left panels; *Alchemilla* sp. and *G. montanum* from six sites) and the second dataset (right panels; *Alchemilla* sp., *D. octopetala* and *G. montanum* from two sites). Only ASVs with an occupancy of 0.25 were tested (131 ASVs and 161 ASVs for dataset 1 and dataset 2, respectively). Different shapes and shape colours show the different tissues and sites, respectively.

**Fig. S6.**
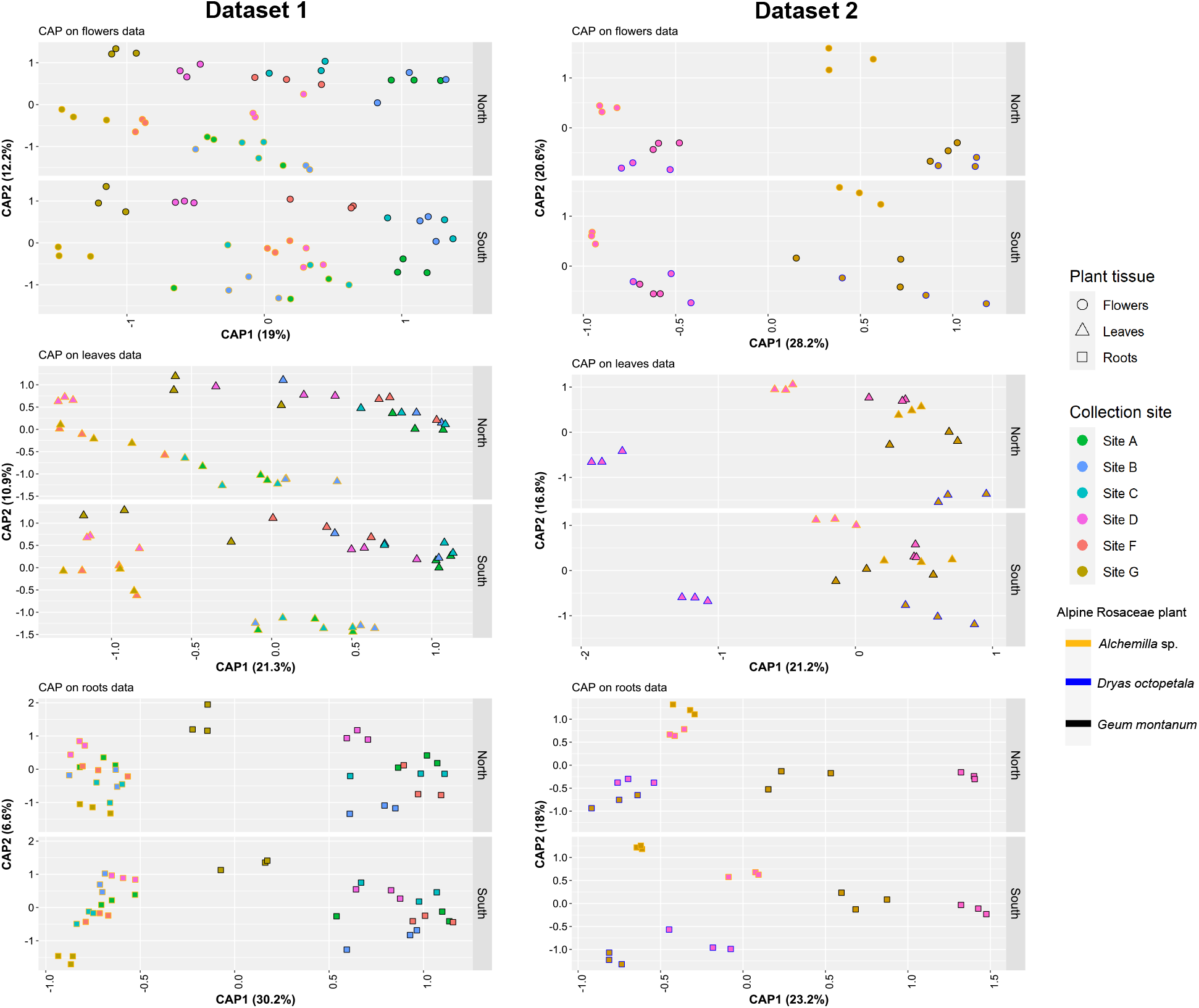
Constrained analysis of principle (CAP) coordinates based on Bray–Curtis dissimilarity matrices in each tissue of different alpine Rosaceae plants, collection sites and exposition. The first dataset (left panels; *Alchemilla* sp. and *G. montanum* from six collection sites) and the second dataset (right panels; *Alchemilla* sp., *D. octopetala* and *G. montanum* from two collection sites) are shown. Different symbols, fill and outline colours represent different tissue, plant species and sites, respectively.

**Fig S7.**
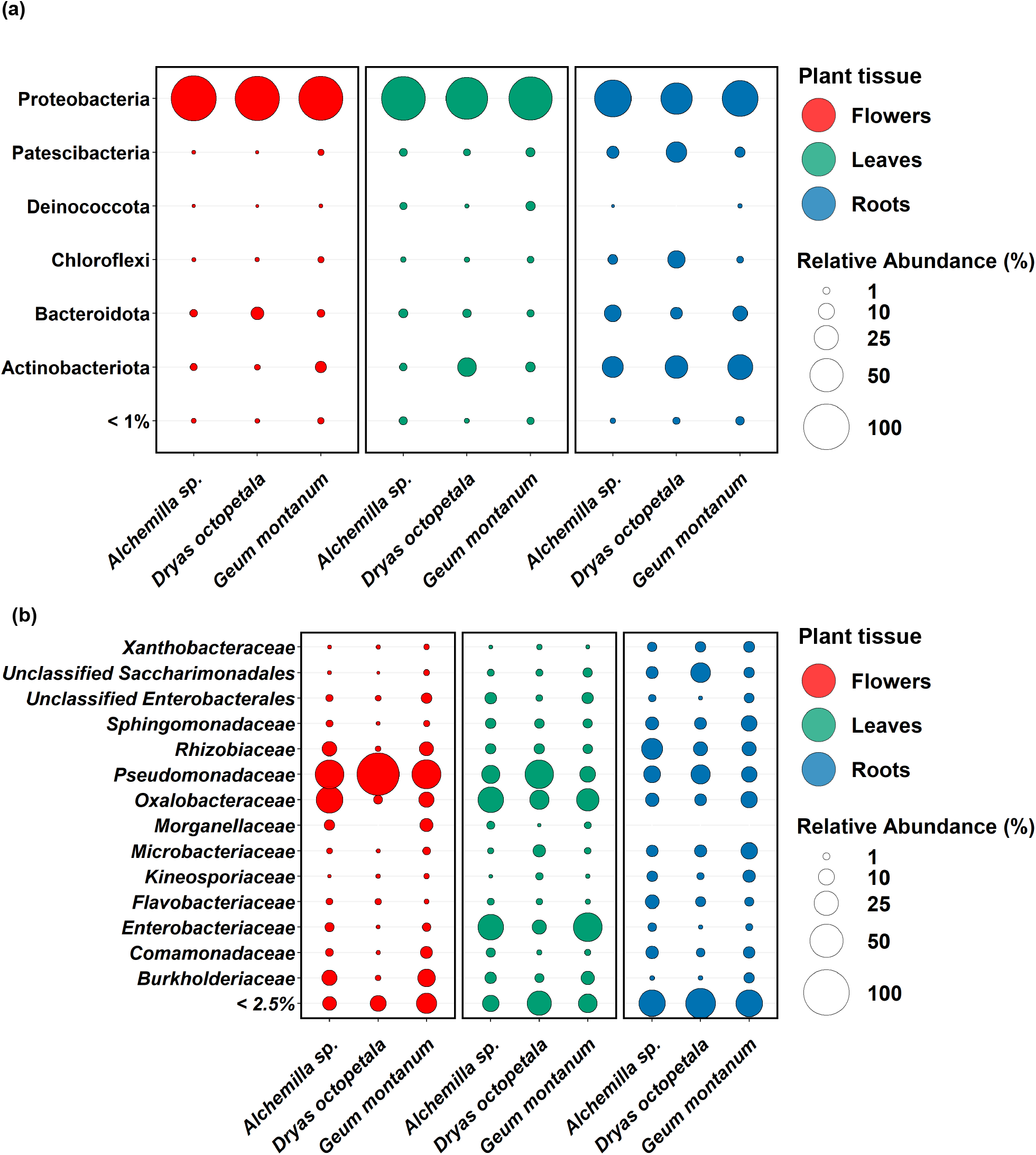
Taxonomic summaries of the endophytic bacterial community structure of alpine Rosaceae plants. Bubble plots of bacterial community classification at the phylum level (a) and family level (b) in different tissues and plant species across all sites and expositions. Only phyla and families with > 1% and > 2.5% mean relative abundance are shown, respectively. Different colours represent different tissues while bubble sizes represent mean relative abundance.

**Fig S8.**
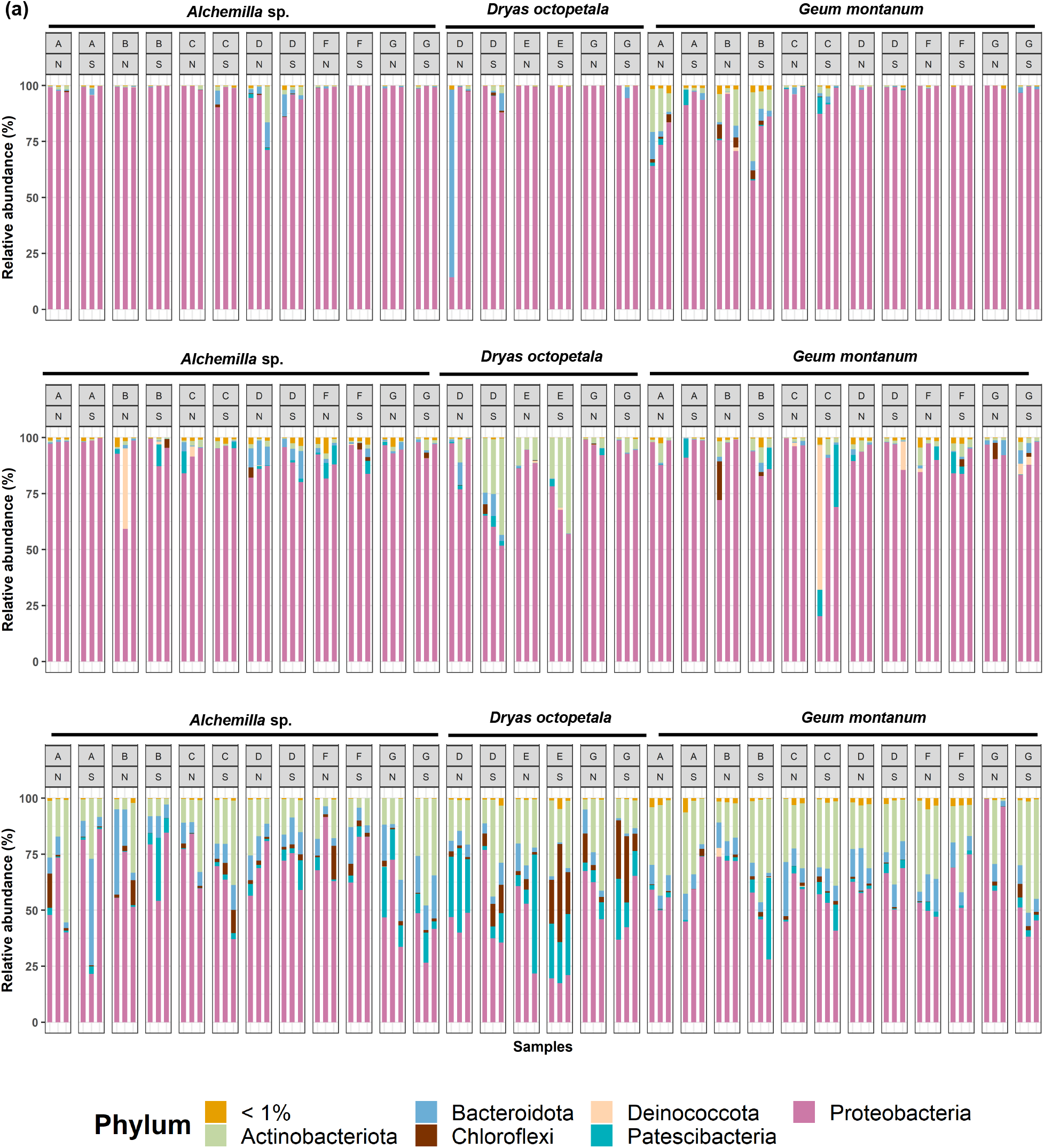
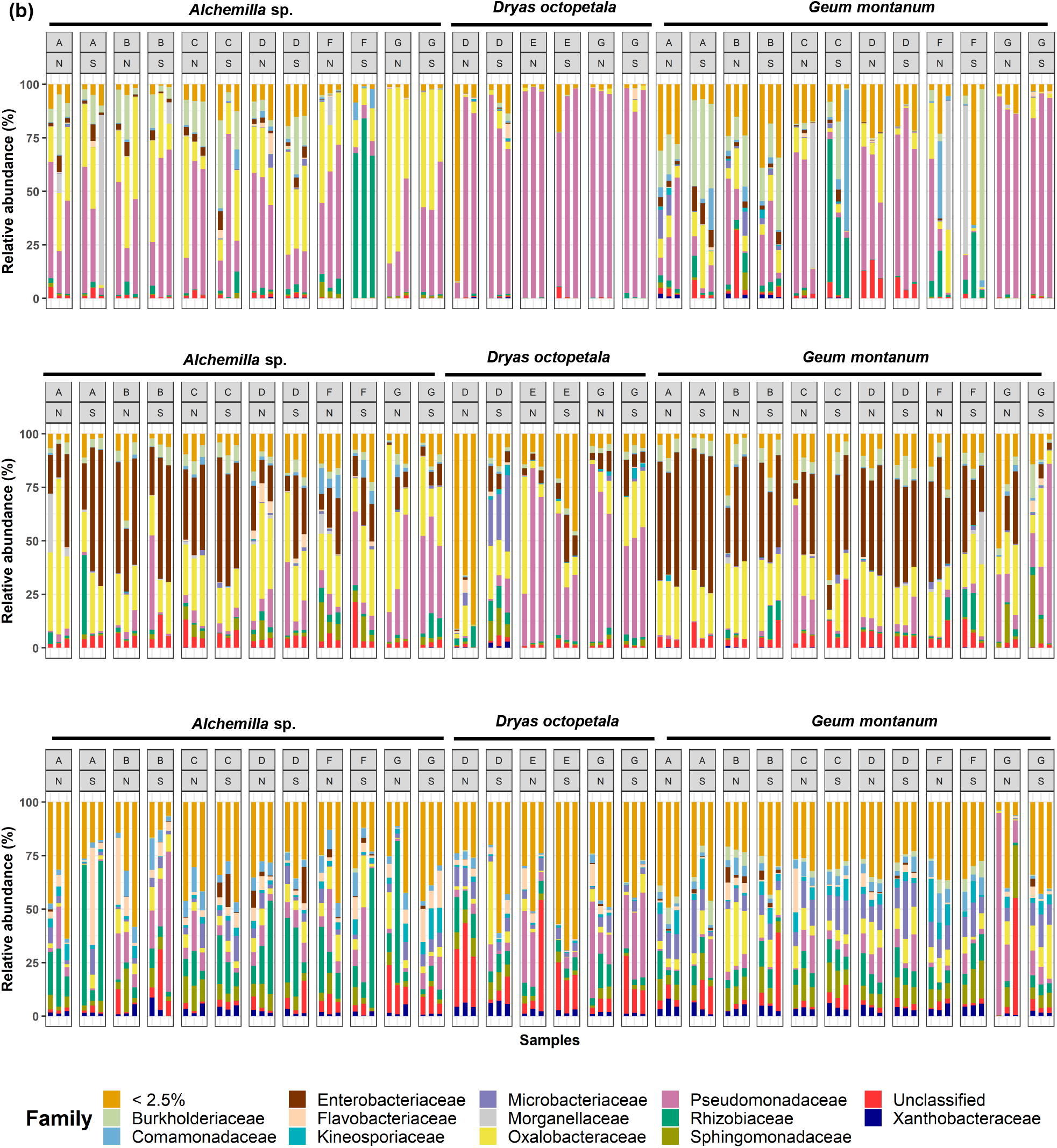
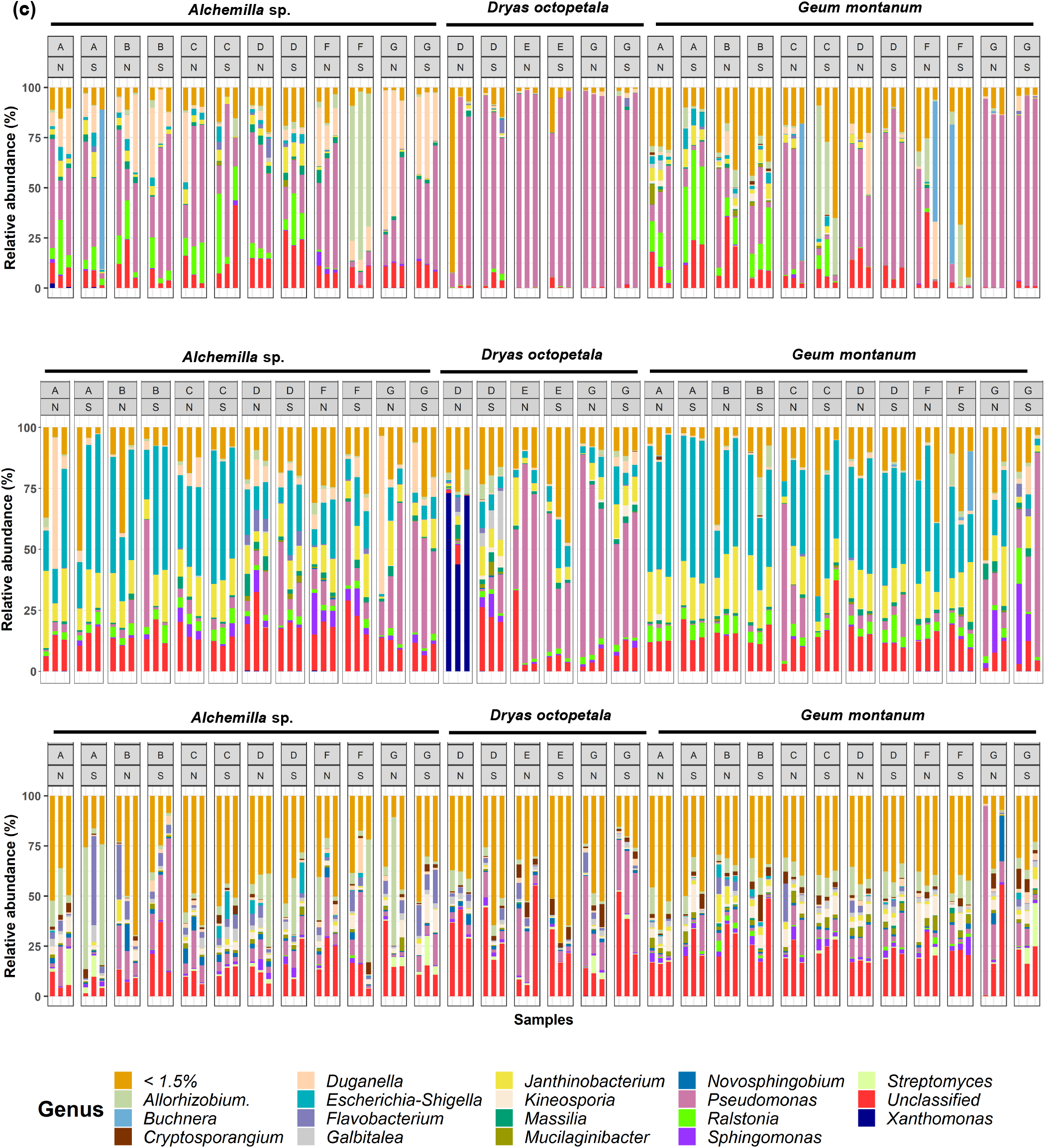
Relative abundance of the endophytic bacterial communities of alpine Rosaceae plants. Bar plots of relative abundance of bacterial communities of *Alchemilla* sp., *Dryas octopetala* and *Geum montanum* at phylum-level (a), family-level (b) and genus-level (c). Bar plots reports data of flowers, leaves and roots for each site (Val di Non, Val di Sole, Val di Pejo, Val Rendena, South Tyrol, Stelvio Park and Val di Fassa; A, B, C, D, E, F and G, respectively) and exposition (N and S) indicated in the grey boxes. Only phyla, families and genera with > 1%, > 2.5% and > 1.5% relative abundance are shown, respectively.

**Fig S9.**
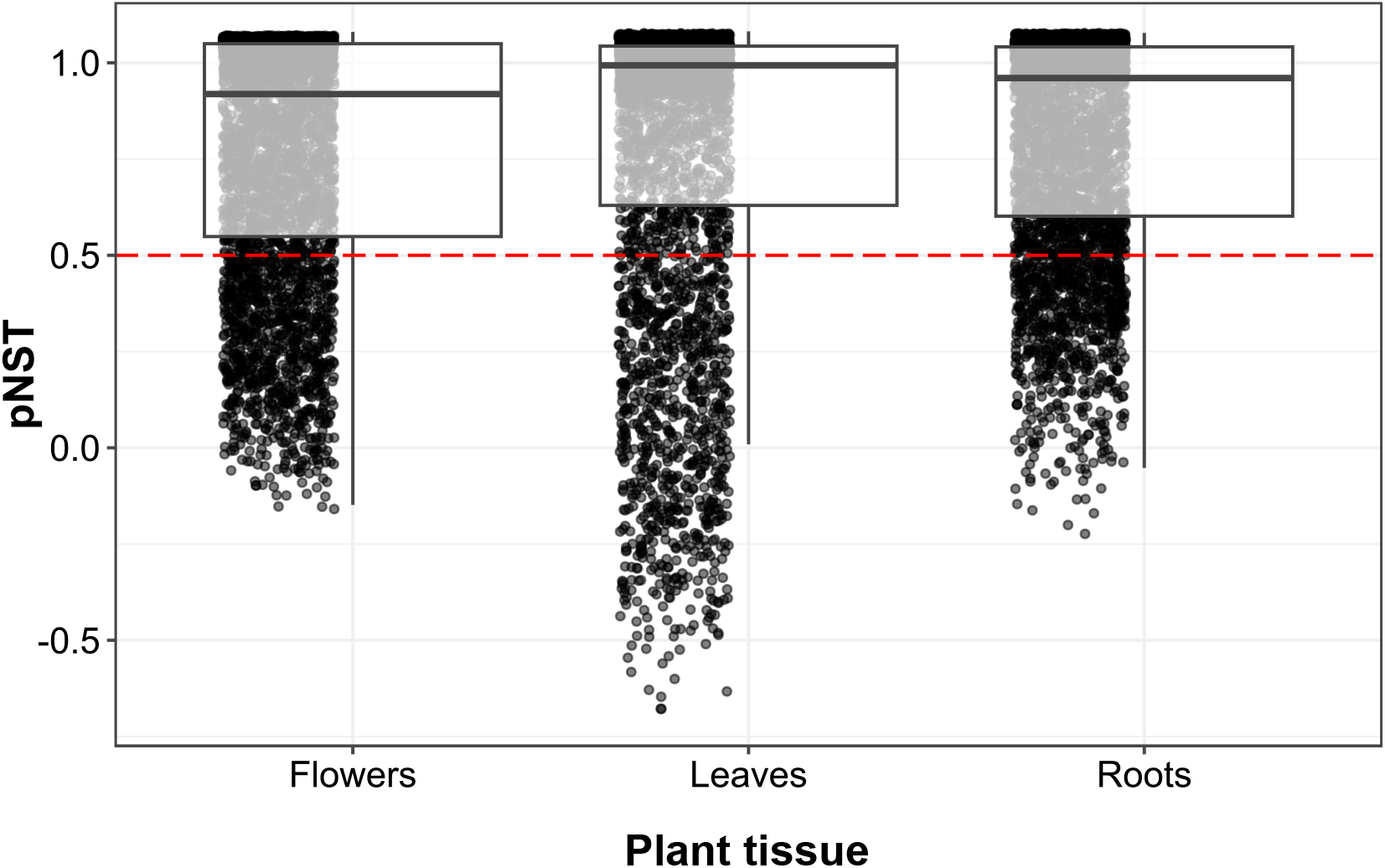
Phylogenetic Normalized Stochasticity Ratios (pNST) for determinism and stochasticity of endophytic bacterial community aggregation in different tissues of alpine Rosaceae plants. The red line indicates the boundary between more deterministic assembly (< 0.5) and more stochastic assembly (> 0.5). Black and grey dots represent each amplicon sequence variant and white squares indicate the median values of 0.75–0.79 in flower, leaf and root tissues.

**Fig S10.**
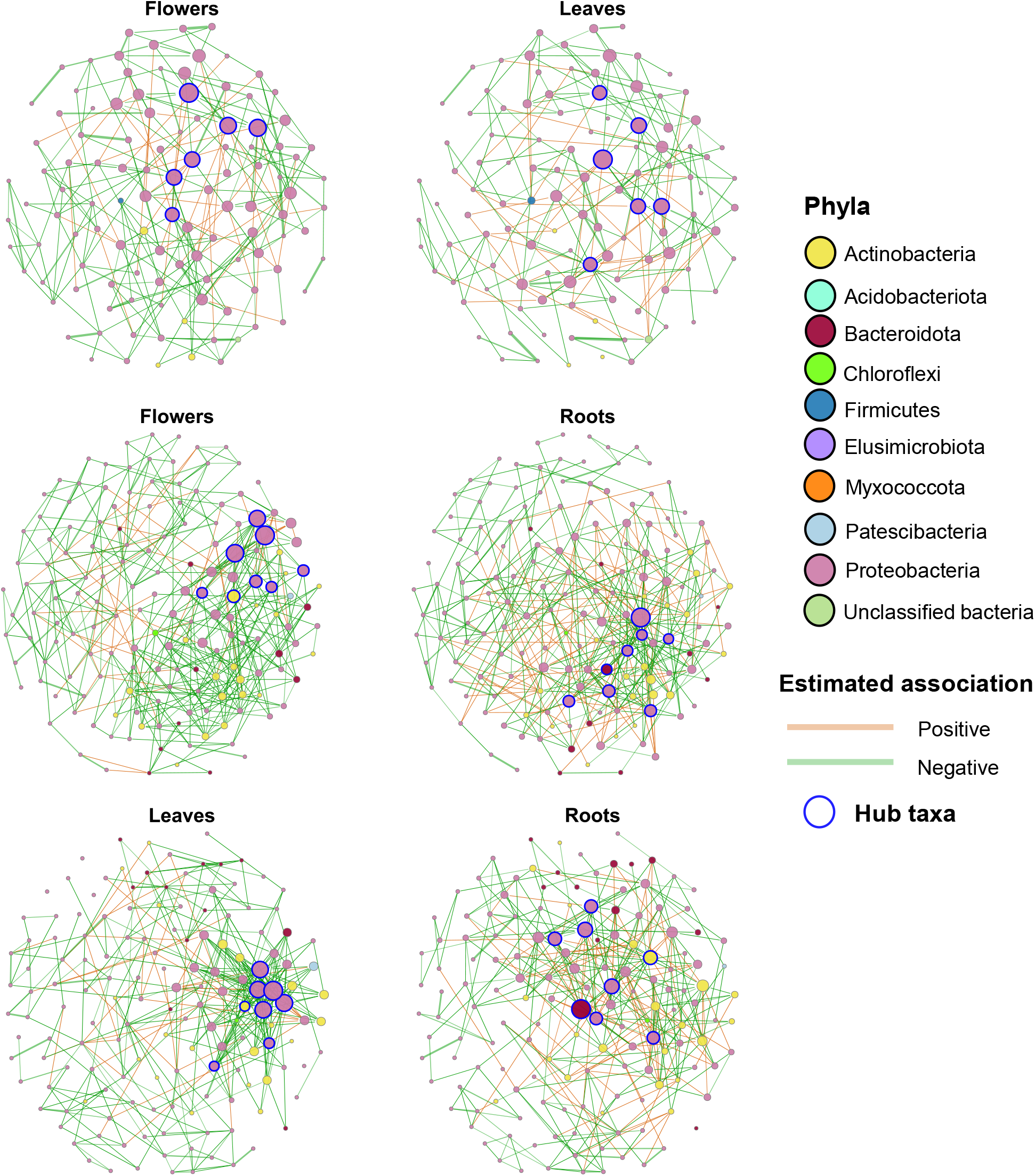
Co-occurrence network comparisons of endophytic bacterial communities alpine Rosaceae plants. Network comparison between flower and root (a) flower and leaf (b) and leaf and root (c) tissues. ASVs with a relative abundance greater than 0.001% and occupancy greater than 0.25 were used for network analysis using the SpiecEasi and NetCoMi R packages. Comparative analysis of networks was achieved between the similarity of most central nodes (i.e. set of ASVs with eigenvector centrality values greater than 95% of the empirical distribution of all eigenvector centralities in the network). Each node corresponds to an ASV, and edges between nodes correspond to either positive (orange) or negative (green) correlations. The thickness of each edge is proportional to the correlation coefficients of the connections. ASVs belonging to different bacterial phyla have distinct colour codes. Node size reflects their eigenvector centrality while nodes with thick blue outlines represent hub taxa.

**Fig S11.**
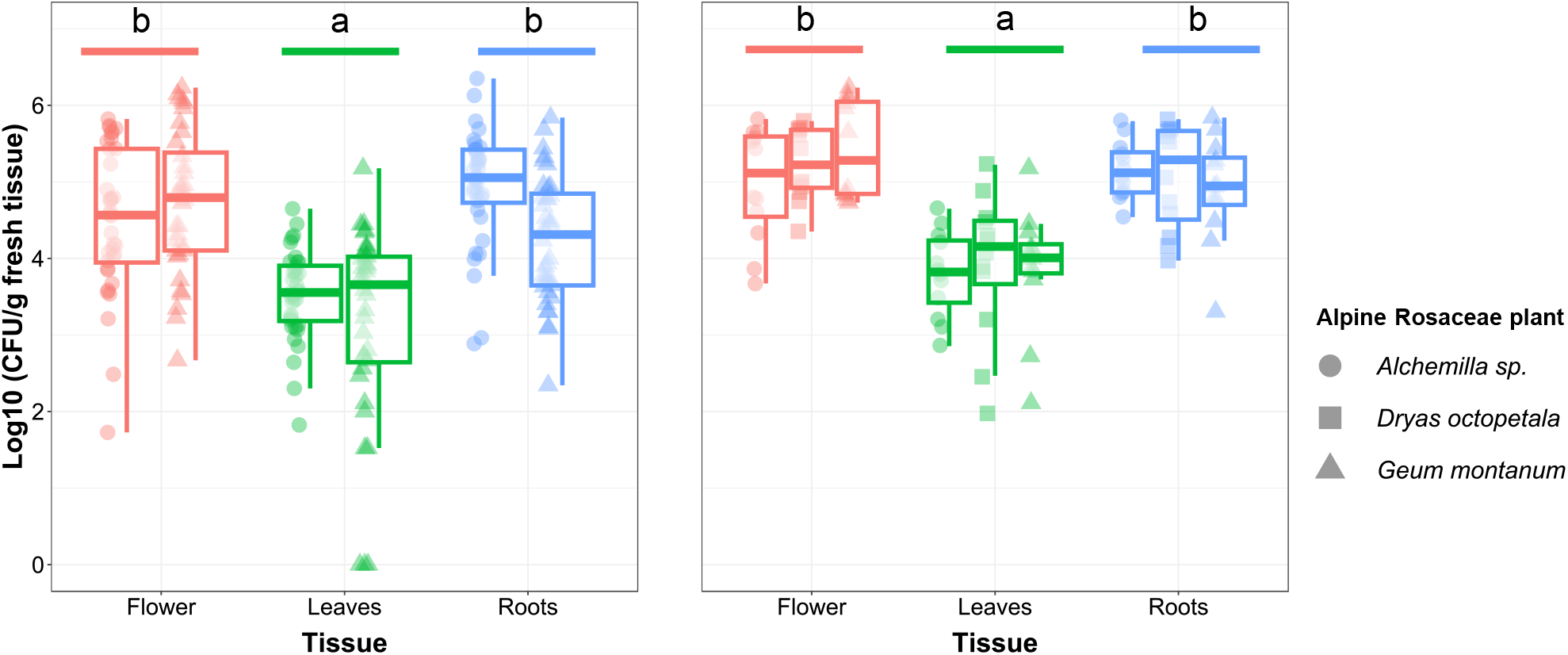
The population density of culturable psychrotolerant bacterial endophytes in alpine Rosaceae plants. Box plots represent log10 colony forming unit (CFU) per gram of fresh weight of flower, leaf and root tissues in the first dataset (a; *Alchemilla* sp. and *G. montanum* from six collection sites) and the second dataset (b; *Alchemilla* sp., *D. octopetala* and *G. montanum* from two collection sites). Values were obtained from three replicates. Different letters indicate significant differences, according to ANOVA followed by post-hoc analysis with estimated marginal mean comparisons (*P* ≤ 0.05).

## Supplementary tables

**Table S1.** Summary of collection site and sample description of alpine Rocaseae plants.

**Table S2.** PCR programs and conditions used for 16S rRNA gene amplicon sequencing (A) and Sanger sequencing (B).

**Table S3.** Summary of the 16S rRNA gene amplicon sequencing analysis of endophytic bacterial communities associated with alpine Rosaceae plants. Raw reads (prior to any filtering steps), total reads (plant and bacterial/archaeal reads), bacterial/archaeal reads (post filtering-out chloroplast and mitochondria reads), observed amplicon sequence variants (richness) and Simpson’s index (alpha-diversity) are reported for each replicate (named 1, 2, and 3) of samples collected form *Alchemilla* sp. (Alc), *Dryas octopetala* (Dry), and *Geum montanum* (Geu) flower (F), leaf (L), and root (R) samples from seven different collection sites (named from A to G) from North (N) or South (S) exposition.

**Table S4.** Bacterial amplicon sequence variants (ASVs) of endophytic bacterial communities associated with alpine Rocaseae plants. Samples included three alpine Rosaceae plants: *Alchemilla* sp. (Alc), *Dryas octopetala* (Dry), and *Geum montanum* (Geu), and three tissues: flowers (F), leaves (L) and roots (R) that were collected from seven collection sites (from A to G) and two expositions: North (N) and South (S). Taxonomic annotation (columns B-H), filtered read counts (columns I-JR) and relative abundances (columns JS-UB) are reported for each replicate sample (from 1 to 3). The mean relative abundance and occupancy of each ASV are shown in columns (UC and UD, respectively). The most abundant ASVs with a mean relative abundance of > 0.1% are highlighted in bold. ASVs classified as core and non-core taxa are shown in column UE. ASVs with no reads or singletons, as well as very low-abundance ASVs with a maximum relative abundance below 0.1% per sample were filtered-out from the ASV table. Core ASVs are indicated in column **UE.**

**Table S5.** Analysis of variance (ANOVA) of linear models (LMs) on bacterial richness for the first dataset (A; *Alchemilla* sp. and *G. montanum* from six collection sites) and second dataset (B; *Alchemilla* sp., *D. octopetala* and *G. montanum* from two collection sites) of samples from alpine Rosaceae plants. Results of the model performance to select the most appropriate model (i.e. model with either fixed effect or with interactions having the lowest root mean squared error (RMSE) for the analysis are shown in the top. A post-hoc analysis with estimated marginal mean (EMM) comparisons was carried-out to better highlight differences between levels in each factor (alpine Rosaceae plant, plant tissue, site and exposition) (*P* ≤ 0.05).

**Table S6.** Analysis of variance (ANOVA) of linear models (LMs) on bacterial alpha-diversity for the first dataset (A; *Alchemilla* sp. and *G. montanum* from six collection sites) and second dataset (B; *Alchemilla* sp., *D. octopetala* and *G. montanum* from two collection sites) of samples from alpine Rosaceae plants. Results of the model performance to select the most appropriate model (i.e. model with either fixed effect or with interactions having the lowest root mean squared error (RMSE)) for the analysis are shown in the top. A post-hoc analysis with estimated marginal mean (EMM) comparisons was carried-out to better highlight differences between levels in each factor (alpine Rosaceae plant, plant tissue, collection site and exposition) (*P* ≤ 0.05).

**Table S7**. Canonical analysis of principal coordinates (CAP) permutation test results for the first dataset (A; *Alchemilla* sp. and *G. montanum* from six collection sites) and second dataset (B; *Alchemilla* sp., *D. octopetala* and *G. montanum* from two collection sites) of samples from alpine Rosaceae plants.

**Table S8.** Factors affecting the beta-diversity of endophytic bacterial communities of alpine Rosaceae plants. Results of a permutational multivariate analysis of variances (PERMANOVA) on bacterial Bray–Curtis dissimilarities for the first dataset (A; *Alchemilla* sp. and *G. montanum* from six collection sites) and second dataset (B; *Alchemilla* sp., *D. octopetala* and *G. montanum* from two collection sites) of samples from alpine Rosaceae plants. The percentage of explained variance (R2 = contribution coefficient) of each factor to the community beta-diversity is reported in columns E and N. Intra-factor pairwise comparisons between levels in each factor (Rosaceae plant, plant tissue, collection site and exposition) was carried-out using the pairwise.perm.manova function from the RVAideMemoire R package with P-value adjustment using Benjamini-Hochberg method (*P* ≤ 0.05). Multivariate homogeneity of group dispersions was performed using the betadisper function from the vegan package, followed by ANOVA. Intra-factor pairwise comparisons between factor levels was carried-out using permutational test (999 iterations) with the permutest function from the vegan R package.

**Table S9.** Factors affecting the beta-diversity of endophytic bacterial communities of alpine Rosaceae plants identified through multivariate generalized linear models (mGLMs). Analysis was performed for the first dataset (A; *Alchemilla* sp. and *G. montanum* from six collection sites) and second dataset (B; *Alchemilla* sp., *D. octopetala* and *G. montanum* from two collection sites) of samples from alpine Rosaceae plants. Rarefied bacterial count data of amplicon sequence variants (ASVs) with 0.25 occupancy were used in this analysis. An analysis of deviance was calculated with a likelihood-ratio test using a permutational test (999 iterations, Monte Carlo resampling).

**Table S10.** Factors affecting the beta-diversity of endophytic bacterial communities in flowers, leaves and roots of alpine Rosaceae plants. Results of a permutational multivariate analysis of variances (PERMANOVA) partitioning test by tissues on bacterial Bray–Curtis dissimilarities for the first dataset (A; *Alchemilla* sp. and *G. montanum* from six collection sites) and second dataset (B; *Alchemilla* sp., *D. octopetala* and *G. montanum* from two collection sites) of samples from alpine Rosaceae plants. Pairwise comparisons between levels within each factor was performed using permutation MANOVAs. The percentage of explained variance (R2 = contribution coefficient) of each factor to the community beta-diversity is reported in columns E, N and W. Host-environment effects index (HEEI = relative contribution of Rosaceae plant/relative contribution of collection site) is reported in columns H, Q and Z.

**Table S11.** Main ecological processes governing the assembly of flower-, leaf- and root-associated communities of alpine Rosaceae plants analysed through Infer Community Assembly Mechanisms by Phylogenetic-bin-based null model (iCAMP). Relative abundances of bins and their top ASVs (A). Relative importance of each bin to the different processes governing community assembly (B). Drift and others (DR), homogenizing dispersal (HD), dispersal limitation (DL), homogeneous selection (HoS), heterogeneous selection (HeS) is reported in column G. The dominant process and its relative importance are shown for each bin.

**Table S12.** Indicator taxon analysis with random forest (RF) machine learning identified bacterial amplicon sequence variants (ASV) based on all the four factors (Rosaceae plant, tissue, site and exposition). Bacterial ASVs based on individual factor with significant mean decrease accuracy from the first dataset (A; *Alchemilla* sp. and *G. montanum* from six collection sites) and second dataset (B; *Alchemilla* sp., *D. octopetala* and *G. montanum* from two collection sites) of samples from alpine Rosaceae plants are reported. P-values for each level within the factor are shown. Taxonomy (columns W and BB) indicates kingdom (k), phylum (p), class (c), order (o), family (f), genus (g), and species (s) of selected ASVs (columns V and BA).

**Table S13.** Significantly differentially abundant amplicon sequence variants (ASV) from the first dataset (A; *Alchemilla* sp. and *G. montanum* from six collection sites) and second dataset (B; *Alchemilla* sp., *D. octopetala* and *G. montanum* from two collection sites) of samples from alpine Rosaceae plants. Log fold change (lfc), standard errors (se) of lfc, W test statistics (W = lfc/se), p-values (obtained from two-sided Z-test using the W test statistic), q values (adjusted p-values obtained by applying Benjamini-Hochberg method) are reported for each ASV in the flower and leaf tissues compared with the root tissue. Columns starts with Tissue_diff_abund represent significant ASVs in the flower or leaf tissues compared with the root tissue.

**Table S14.** Summary of network properties of endophytic bacterial co-occurrence networks of Rosaceae plants. Networks for each tissue (flower, leaf and root) and network comparison between them (flower vs root, flower vs leaf and leaf vs root) are reported. Hub nodes were identified based on eigenvector centrality values. The Jaccard index was used for assessing how different the sets of most central nodes are between the two networks (0 if the sets are completely different and 1 for exactly equal sets).

**Table S15.** Taxonomic annotation at the genus level for the culturable psychrotolerant bacterial endophytes from alpine Rosaceae plants. Blast results of the top ten hits of the bacterial culture collection against the NCBI 16S ribosomal RNA database are reported. NCBI accession numbers for our isolate (column B) and the closest relative to our isolate (column C), identity percentage and alignment length are presented.

**Table S16.** Analysis of variance (ANOVA) of linear models (LMs) on the population density of culturable psychrotolerant bacterial endophytes for the first dataset (A; *Alchemilla* sp. and *G. montanum* from six collection sites) and second dataset (B; *Alchemilla* sp., *D. octopetala* and *G. montanum* from two collection sites) of samples from alpine Rosaceae plants. Results of the model performance to select the most appropriate model (i.e. model with either fixed effect or with interactions having the lowest root mean squared error (RMSE)) for the analysis are shown in the top. A post-hoc analysis with estimated marginal mean (EMM) comparisons was carried-out to better highlight differences between levels in each factor (alpine Rosaceae plant, plant tissue, site and exposition) (*P* ≤ 0.05).

**Table S17.** Identification of representative psychrotolerant bacterial endophytes of Rosaceae plants. Blast results of Sanger sequences from the culturable psychrotolerant bacterial endophytes against all recovered bacterial amplicon sequence variants (ASV) from this study are shown. Only the top ten hits are shown for each culturable isolate with the identity percentage and alignment length against an ASV.

